# Neuronal Aralar/AGC1 deficiency impairs social behavior through dopaminergic and redox dysfunction and is rescued by β-hydroxybutyrate

**DOI:** 10.64898/2026.11.12.766132

**Authors:** Eduardo Herrada-Soler, Nerea Murugarren, Miguel Angel Serrano-Lope, María José Casarejos, Javier Palazuelos, Jorgina Satrústegui, Araceli del Arco, Beatriz Pardo

## Abstract

**Background:** Pathogenic variants in SLC25A12/AGC1 (Aralar) have been linked to autism spectrum disorder (ASD), yet the mechanisms connecting neuronal metabolic dysfunction to altered social behavior remain poorly understood. Aralar is a key component of the malate-aspartate shuttle, supporting neuronal oxidative metabolism and lactate utilization. Here, we investigated the consequences of postnatal neuronal Aralar deletion and the therapeutic potential of β-hydroxybutyrate (BHB).

**Methods:** We generated neuron-specific Aralar knockout mice (Neu Aralar-KO) using CamKIIα-Cre, producing efficient recombination in cortex, hippocampus, and striatum. Behavioral, transcriptomic, neurochemical, morphometric, histological, and biochemical analyses were performed in adult mice. The effects of systemic BHB administration were assessed using behavioral, metabolic, oxidative stress, and transcriptomic endpoints.

**Results:** Despite a 55-65% reduction of Aralar expression in forebrain regions, Neu Aralar-KO mice displayed normal growth, survival, brain structure, and myelination. Unexpectedly, loss of neuronal Aralar did not impair spatial memory, episodic memory, or reversal learning. In contrast, Neu Aralar-KO mice exhibited hyperactivity and a marked deficit in sociability, revealing an ASD-relevant behavioral phenotype. Morphological analysis showed significant remodeling of striatal dendritic spines without changes in spine density. Transcriptomic profiling identified 433 differentially expressed genes, the vast majority being upregulated, with enrichment of pathways related to neuropeptide signaling, synaptic transmission, membrane excitability, and behavior. Notably, genes involved in dopamine (DA) and glutamate handling, synaptic connectivity, neuromodulatory signaling, and ASD-associated pathways were strongly induced. Neurochemical analyses revealed preserved glutamate and GABA levels but reduced striatal DA content accompanied by an increased DOPAC/DA ratio, indicating enhanced DA degradation. DAergic neurons of the ventral tegmental area (VTA) showed minimal recombination, suggesting that DA dysfunction arises through non-cell-autonomous mechanisms. Consistent with increased DA oxidative catabolism, Neu Aralar-KO striata displayed elevated lipid peroxidation, induction of glutathione-related pathways, upregulation of antioxidant genes, and astrocyte activation. Remarkably, BHB treatment restored sociability, normalized striatal metabolic alterations, reduced oxidative stress, and abolished genotype-dependent neurochemical abnormalities. Transcriptomic analyses further revealed that Aralar deficiency recapitulates a substantial fraction of the fasting/BHB-induced epigenetic transcriptional program.

**Conclusions:** Postnatal neuronal Aralar deficiency uncouples social behavior from cognitive performance and drives ASD-like phenotypes through a mechanism linking neuronal metabolic dysfunction, oxidative stress, glial activation, and impaired DAergic homeostasis. The rescue of behavioral, metabolic, and redox abnormalities by BHB identifies restoration of metabolic homeostasis as a major therapeutic mechanism and highlights metabolic interventions as promising strategies for AGC1 deficiency and ASD-associated disorders involving neuronal metabolic dysfunction.

## INTRODUCTION

Aralar/AGC1, the mitochondrial aspartate-glutamate carrier, constitutes the regulatory component of the NADH malate-aspartate shuttle (MAS) (Palmieri et al., 2001; Satrustegui et al., 2007). Aralar/AGC1 is a calcium-sensitive carrier that contains EF-hand calcium-binding motifs, enabling MAS stimulation and activation of mitochondrial metabolism by cytosolic calcium (Palmieri et al., 2001; Pardo et al., 2006; Llorente-Folch et al., 2013a; Perez-Liébana et al., 2022). Aralar deficiency in humans causes the rare disorder known as “early infantile epileptic encephalopathy” (OMIM # 612949), characterized by neurodevelopmental arrest, severe hypomyelination, hypotonia, and refractory seizures (Wibom et al., 2009; Pardo et al., 2022, for review). Global Aralar KO mice die prematurely and exhibit key features that parallel the human syndrome, including neurodevelopmental delay, hypomyelination, hypotonia, and epilepsy (Jalil et al., 2005). Thus, the Aralar-KO mouse provides a robust model that faithfully recapitulates the pathophenotype observed in AGC1-deficient patients. Aralar is predominantly expressed in excitable tissues (del Arco and Satrustegui, 1998) but also in kidney, in blood and immune cells. Within the brain, Aralar expression has been mainly shown in neurons; with no evidence of Aralar protein in mouse oligodendrocytes (OL) *in vivo* (Ramos et al., 2003; Pardo et al., 2011; Juaristi et al., 2017). However, transcriptomics revealed that OPCs and OLs do express Aralar mRNA but at lower levels than neurons (Cahoy et al., 2008). Within this framework, the contribution of specific Aralar/AGC1 expressing cells to the drastic global phenotype in humans and mice remains unknown.

Neurons, the excitable brain cells with the highest levels of Aralar expression, display unique properties that make them the most energy-demanding cell type in humans (Cheng et al., 2022). It is well established that neurons rely predominantly on mitochondrial OXPHOS for ATP production and that they are particularly vulnerable to mitochondrial dysfunction. These considerations led us to hypothesize that the selective loss of Aralar in neurons is the major driver of the severe phenotype found in AGC1 deficiency. Although the absence of neuronal Aralar primarily affects neuronal physiology, it may also disrupt brain functions depending on glial cells, such as astrocytic glutamine (Gln) synthesis or oligodendrocyte-mediated myelin formation. Glial dysfunctions in AGC1 deficiency likely stem from reduced availability of neuron-derived metabolites. Specifically, diminished neuronal-born N-acetylaspartate (NAA) is proposed to underlay the characteristic hypomyelination, while deficits in neuronal Aspartate (Asp) are likely to contribute to impaired astrocytic Glutamate (Glu)/Gln synthesis observed in AGC1 deficiency (Jalil et al., 2005; Wibom et al., 2009; Pardo et al., 2011).

Notably, AGC1/Aralar has been shown to be essential in neurons for: a) the synthesis of Asp and NAA (Jalil et al., 2005; Llorente-Folch et al., 2013b); b) the utilization of lactate as an energy substrate (Pardo et al., 2006; Llorente-Folch et al., 2016); c) the control of mitochondrial respiration supported by glucose (Llorente-Folch et al., 2013a, Pérez-Liébana et al., 2020, 2022); d) electrogenesis (Gómez-Galán et al., 2011); and e) the synthesis/metabolism of neurotransmitters such as Glu and dopamine (DA) (Pardo et al., 2011; Llorente-Folch et al., 2013b), among other functions. In neurons, but not in astrocytes, Aralar-deficiency impairs basal and stimulus-evoked mitochondrial respiration fueled by glucose. It also restricts glycolytic flux as well as the production and mitochondrial utilization of pyruvate (Gómez-Galán et al., 2011; Llorente-Folch et al., 2013a; Juaristi et al., 2017; Pérez-Liébana et al., 2020, 2022). Aralar/AGC1-MAS is also required to couple neuronal activity to bioenergetic responses, enabling neurons to cope with stimuli such as NMDA. Importantly, failure of AGC1/Aralar-MAS prevents neuronal lactate oxidation, abolishing lactate’s protective effects against excitotoxicity (Llorente-Folch et al., 2016). These deficits may constitute the mechanistic basis for alterations in neuronal excitability and may contribute to impairments in learning and memory processes, among other consequences.

Interestingly, a substantial body of evidence links impairments in the Aralar/AGC1 to autism spectrum disorder (ASD). Both polymorphisms in the *SLC25A12* gene and altered levels of the Aralar/AGC1 protein have been reported in individuals with ASD (Ramoz et al., 2004; Segurado et al., 2005; Kim et al., 2011; Aoki and Cortese, 2016; Ghavi et al., 2024). Regarding social behavior, AGC1-deficient individuals exhibit reduced eye contact and impaired communication (Wibom et al., 2009), whereas global Aralar-KO mice display behavioral deficits in homing and in home-cage social interaction (Llorente-Folch et al., 2013b). ASD is a complex neurodevelopmental condition characterized by impairments in social interaction and communication, along with restricted and stereotyped behavioral patterns. Common comorbidities include hyperactivity and a high incidence of epileptic activity. Indeed, individuals with AGC1 deficiency present treatment-refractory epilepsy and global Aralar-KO mice exhibit hyperactivity with spontaneous seizures.

The pathogenic pathways implicated in ASDs involve defects in synaptic function (Zoghbi et al., 2003; Persico and Bourgeron, 2006), pointing to either excessive or diminished excitatory synaptic connectivity (Chao et al., 2007; Hanson and Madison, 2007) and changes in dendritic spine density and structure (Chidambaram et al., 2019). Additionally, the social motivation hypothesis of autism posits that children with ASD do not perceive social stimuli as inherently rewarding. At this respect, multiple lines of evidence link dopaminergic (DAergic) modulation of the mesolimbic pathway to the processing of social reward (Scott-Van Zeeland et al., 2010; Lin et al., 2012). Human genetic studies have identified contributions of DA-related genes to the modulation of social behavior, and mutations affecting components of the mesolimbic DAergic pathway have been associated with ASD (Groppe et al., 2013; Enter et al., 2012; Eisenegger et al., 2013; Staal, 2015; Bowton et al., 2014). Altogether, these findings point to deficits in social-reward processing in ASD, accompanied by genetic and functional abnormalities in the mesolimbic DA system. In this regard, the striatal DAergic neurons involved in motor control and social reward are primarily affected in global Aralar-KO mice (Llorente-Folch et al., 2013b; Pérez-Liébana et al., 2020). Given the high prevalence of ASD (1 in 54 children; Maenner et al., 2020) and its profound impact, it is critical to investigate its mechanistic etiology and to develop effective therapies. For this reason, we aim to elucidate whether dysfunction of the Aralar/AGC1–MAS pathway may play a role in ASD, and, if so, to identify the underlying mechanisms and potential therapeutic strategies.

Our aim is to understand the etiological basis of the phenotype observed in global Aralar/AGC1 deficiency and to determine the specific contribution of distinct brain cell types, particularly neurons. We hypothesized that neuron-specific Aralar ablation may be sufficient to recapitulate a phenotype closely resembling that of the global Aralar-KO mice. The severe phenotype and premature death of the global KO animals before completion of neurodevelopment preclude further investigations into the molecular mechanisms underlying the pathology. To dissect the neuronal function of ARALAR-MAS, we generated a conditional neuronal Aralar knockout (Neu Aralar-KO) mice. For this purpose, (*Aralar^fl/fl^*) mice were crossed with mice expressing Cre recombinase under the control of the neuron-specific calcium-calmodulin protein kinase-II (*CamkIIα*) promoter. The present study includes comprehensive general, phenotypic and behavioral characterization of the Neu Aralar-KO mice, together with transcriptomic and metabolomic analyses aimed at identifying key molecular players involved in the observed deficits. Surprisingly, Neu Aralar-KO mice exhibit preserved life expectancy, normal gross brain parameters with intact brain structure and postnatal myelination when compared with littermate controls. Moreover, contrary to expectations, Neu Aralar-KO animals do not display learning-memory impairments. But, interestingly, Aralar Ablation in neurons do generate a specific deficiency in social behavior and hyperactivity; associated with increased striatal oxidative stress, impairments in DAergic metabolism, and alterations in dendritic spine morphology. Notably, chronic BHB treatment successfully rescued the social deficits, oxidative stress, and metabolic impairments observed in the striatum of adult Neu Aralar-KO mice. This result highlights the feasibility of BHB, the most abundant ketone body produced during the ketogenic diet, as a therapeutic strategy even under a non–carbohydrate-restricted dietary regimen.

## RESULTS

### 1. Postnatal disruption of Aralar in neurons from cortex, hippocampus and striatum does not cause changes in growth, survival or myelination

To generate neuron-specific *Aralar* knock out mice, referred to here as Neu Aralar-KO, mice harboring loxP sites flanking exon 3 (Aralar^fl/fl^) (García-Lobo et al., 2026), were mated with a transgenic line expressing Cre recombinase under the control of the neuron-specific calcium-calmodulin protein kinase-II*α* (*CamkIIα*) promoter and carrying Ai6, a Cre-responsive reporter cassette expressing the fluorescent protein ZsGreen (CaMKIIα-Cre; Rosa:Ai6 mouse, Figure 1A, 1B). Offspring from these crosses were genotyped to selected Neu Aralar-KO (*Aralar*^fl/fl^; *CaMKIIα-Cre^+/-^*; *Rosa:AiC^+/-^)* and control (*Aralar*^+/+^; *CaMKIIα-Cre^+/-^*; *Rosa:AiC^+/-^)* mice (Figure 1B, 1C).

**Figure 1.**
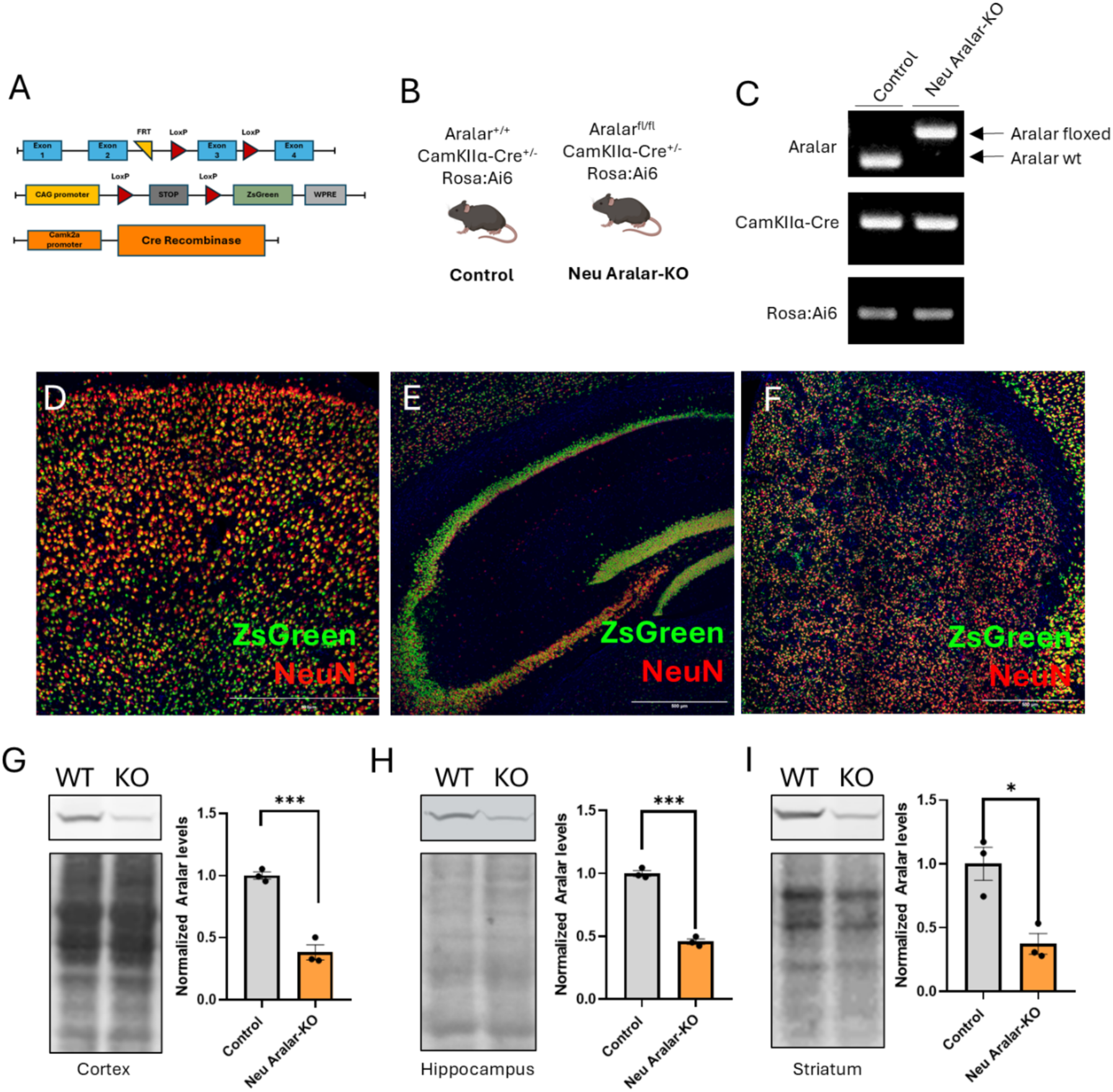
Generation of Neu Aralar-KO mice and validation of neuronal recombination and Aralar ablation across brain regions. **(A)** Schematic representation of the Aralar^fl/fl^, CamKIIα-Cre and Rosa:Ai6 alleles present in (**B**) Control (Aralar^+/+^; CamKIIα-Cre^+/−^; Rosa:Ai6) and littermates Neu Aralar-KO (Aralar^fl/fl^; CamKIIα-Cre^+/−^; Rosa:Ai6) mice. (**C**) Representative PCR genotyping of Control and Neu Aralar-KO animals, showing the floxed and wild-type *Slc25a12* (Aralar) alleles, the CamKIIα-Cre transgene and the Rosa:Ai6 reporter. Representative images of neuronal recombination, visualised as co-localisation of the ZsGreen reporter (green) with NeuN immunostaining (red) and DAPI (blue), in cortex (**D**), hippocampus (**E**) and striatum (**F**). Representative immunoblots and quantification of Aralar protein levels in cortex (**G**), hippocampus (**H**) and striatum (**I**) of Control and Neu Aralar-KO mice at 7 months of age; Ponceau total protein staining was used as loading control. Data are presented as mean ± SEM (n = 3 mice per group). Statistical analysis was performed by unpaired Student’s *t*-test; *p ≤ 0.05, ***p ≤ 0.001. Scale bars, 500µm (D–F).

Recombination efficiency was assessed using ZsGreen reporter and immunofluorescence staining of brain sections with the neuronal marker NeuN. In the cortex, approximately 75% of NeuN-positive neurons co-expressed ZsGreen, compared with 70% in the hippocampus and up to 90% in the striatum (Figure 1D-F), indicating a robust neuronal recombination within these regions. No recombination was found in the cerebellum (data not shown), consistent with the known expression pattern of CaMKII (Minichiello et al., 1999; Henry and Hohmann, 2012). Aralar protein levels were subsequently analyzed at different postnatal ages (from 2-7 months), in regions exhibiting high CaMKIIα expression. In the cortex (Figure S1A) of Neu Aralar-KO mice, Aralar protein levels remained above 60% of control values between 2 and 3 months of age and declined further from 4 months onwards. By 7 months, ARALAR levels were reduced by 55-65% in the cortex, hippocampus and striatum of Neu Aralar-KO mice relative to controls, with no further changes with age (Figure 1G-I).

Unlike the global Aralar KO mouse or AGC1-deficient patients (Jalil et al., 2005; Wibom et al., 2009; Pardo et al., 2022 for review), Neu Aralar-KO mice had no changes in growth or survival compared to controls (Figure S1B) and display no overt phenotypic abnormalities under standard housing conditions. Similarly, no apparent alterations in brain structures such as cortex, hippocampus or striatum, were observed, nor apparent changes in neuronal numbers (NeuN-positive cells). We did not find any change in ventricle size confirmed by in vivo MRI (magnetic resonance imaging, Figure S1C, D); nor in the extent of myelination, assessed by the analysis of myelin markers such as MBP and MAG (Figure S1E-J). This clearly differs with the finding of enlarged ventricles and hypomyelination observed in the global KO mouse (Jalil et al., 2005; Ramos et al., 2011; Llorente-Folch et al., 2013b) and in AGC1 deficient patients (Wibom et al., 2009).

### 2. Neu Aralar KO mice have preserved memory but display ASD-like behaviors

To investigate whether loss of Aralar specifically in neurons beyond postnatal development might affect behavior or cognition, a battery of behavioral tests was conducted in 7-9 month-old Neu Aralar-KO mice (Figure 2). We first evaluated exploratory activity and anxiety-like behavior in the Open field test (Kraeuter et al., 2019a). In this test, Neu bAralar-KO mice exhibited significant increases in total distance travelled, number of center entries, line crossings and average speed compared to controls (Figure 2A-F), with no changes in time spent in the center quadrant (not shown), suggesting hyperactivity and/or enhanced exploratory behavior.

**Figure 2.**
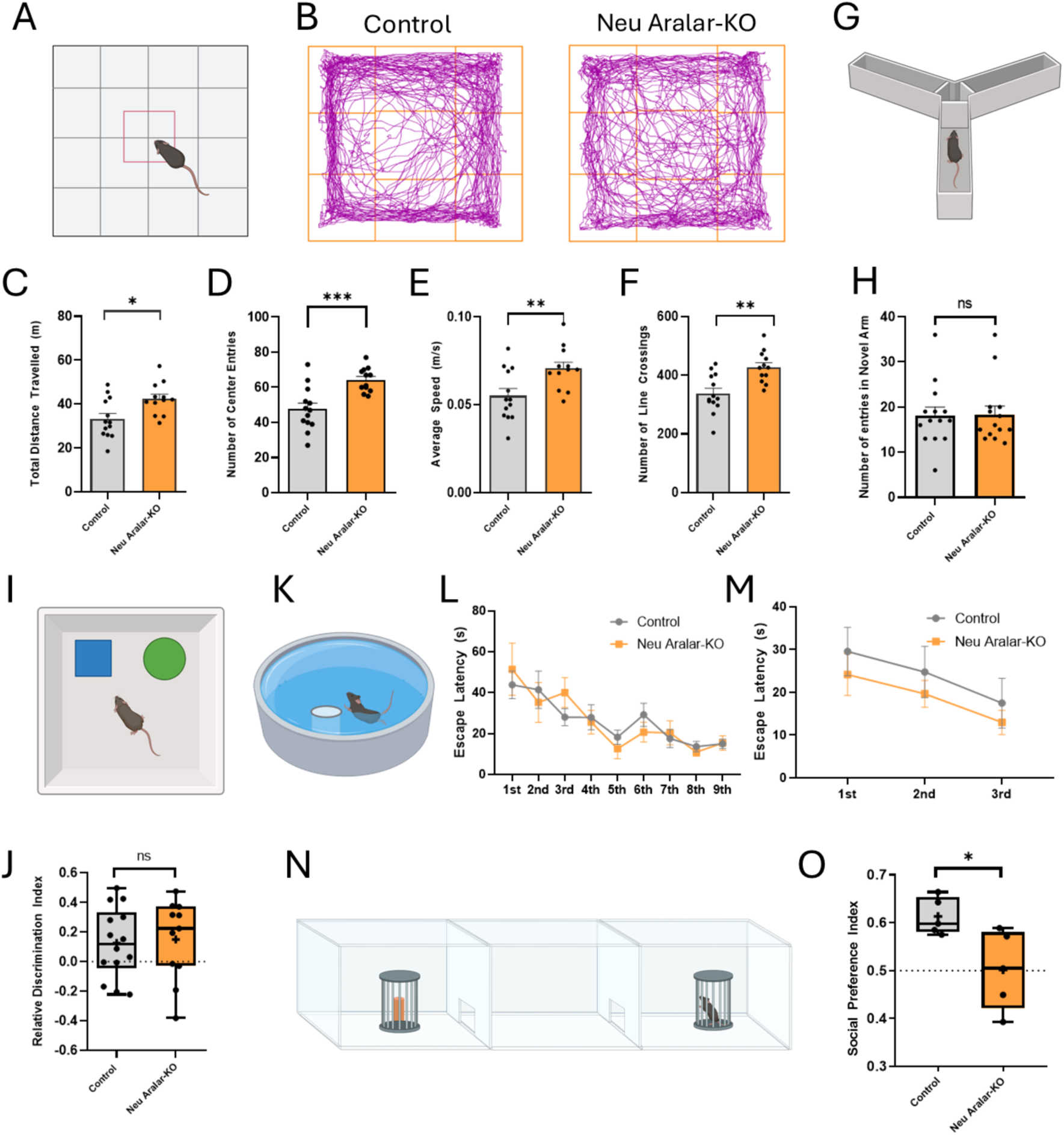
Neuronal Aralar deficiency induces ASD-like social impairment and mild hyperactivity while preserving learning and memory. (**A**) Schematic representation of the open field arena. (**B**) Representative locomotor traces of Control and Neu Aralar-KO mice during the open field test. Ǫuantification of (**C**) total distance travelled, (**D**) number of centre entries, (**E**) average speed and (**F**) number of line crossings (n = 13 Control, n = 12 Neu Aralar-KO). (**G**) Schematic representation of the Y-maze. (**H**) Number of entries into the novel arm during the test phase (n = 14 per group). (**I**) Schematic representation of the novel object recognition test. (**J**) Relative discrimination index between the novel and the familiar object; mean values are represented by “+” (n = 14 Control, n = 11 Neu Aralar-KO). (**K**) Schematic representation of the Morris water maze. (**L**) Escape latency (time from release until the platform is reached) across consecutive acquisition trials and (**M**) escape latency across consecutive reversal-learning trials (n = 13 Control, n = 9 Neu Aralar-KO); L and M were analysed by two-way repeated-measures ANOVA. (**N**) Schematic representation of the three-chamber social interaction test. (**O**) Social preference index, calculated as the time spent sniffing the social stimulus (conspecific) relative to the inanimate object; mean values are represented by “+” (n = 5 per genotype). Data are presented as mean ± SEM. Unless otherwise indicated, statistical analysis was performed using unpaired Student’s *t*-test. n.s., not significant; *p ≤ 0.05, **p ≤ 0.01, ***p ≤ 0.001.

Because the use of lactate in neurons plays a critical role in memory formation (Suzuki et al., 2011; reviewed in Magistretti and Allaman, 2018) and AGC1 deficiency imposes a metabolic impairment of lactate utilization by neurons (Pardo et al., 2006; Llorente-Folch et al., 2016), we evaluated cognitive performance in Neu Aralar-KO mice across a battery of behavioral paradigms (Y-maze, novel object recognition and Morris water maze tests). Spatial memory was assessed using the Y-maze test (Kraeuter et al., 2019b) at both short-term (2 hours, not shown) and long-term (24 hours, Figure 2G and H) intervals. No significant differences between Neu Aralar-KO and controls were observed in any of the evaluated parameters (number of entries in novel arm (Figure 2H); time spent and total distance travelled (data not shown)), indicating intact spatial memory following neuronal Aralar loss. Episodic memory was further examined using the Novel Object Recognition test (Leger et al., 2013; Figure 2I and 2J). Neu Aralar-KO mice showed no impairment in discriminating a novel from a familiar object as evidenced by discrimination indices (DI) comparable to controls (Figure 2J). Multiple object pairs were tested to control for possible bias, yielding consistent results. Neither total distance travelled nor exploration time differed between genotypes during training or testing (not shown). Cognitive flexibility was further evaluated in Neu Aralar-KO mice using the Morris water maze, assessing both spatial learning and reversal learning capabilities (Vorhees et al., 2006; Figure K). Neu Aralar-KO and control mice exhibited similar escape latencies during the acquisition trials, indicating normal learning and memory consolidation (Figure 2L). Following relocation of the hidden platform, both groups adapted equally efficiently to the new platform position, with no genotype-dependent differences in cognitive flexibility measured by reversal learning performance (Figure 2M). Thus, despite the metabolic defect caused by the neuronal Aralar deficiency, spatial memory and cognitive flexibility remained remarkably preserved. Together with the results obtained in other behavioral paradigms, these findings reveal an unexpected resilience of mnemonic processes in Neu Aralar-KO mice.

Given the reported association between *SLC25A12/Aralar* and ASD in humans (Ramoz et al., 2004; Segurado et al., 2005; Kim et al., 2011; Aoki and Cortese, 2016; Ghavi et al., 2024), we assessed social behavior in Neu Aralar-KO mice using the Three-Chamber Social Interaction test, a well-established assay for detecting deficits in sociability in mouse models of ASD (Yang et al., 2011; Figure 2N,2O). This test examines two key components of social behavior: sociability (social reward) and social memory, measured as preference for social novelty. While the social memory test evaluates the ability to discriminate between a familiar and a novel mouse (Yang et al., 2011), it is generally considered less directly related to core ASD-like phenotypes than sociability (Yang et al., 2011). In the sociability test, Neu Aralar-KO mice failed to display the expected preference for a social stimulus over an inanimate object, exhibiting a social preference index close to chance level (≈0.5; Figure 2O). By comparison, control mice showed a robust preference for social interaction, with DI exceeding 0.62. These findings reveal a marked reduction in sociability in Neu Aralar-KO mice, indicating that neuronal Aralar deficiency impairs a core aspect of social behavior relevant to ASD-like phenotypes.

In summary, while Aralar-deficient neurons show reduced capacity to utilize lactate (Llorente-Folch et al., 2016), Neu Aralar-KO mice did not exhibit deficits in episodic and spatial memories or cognitive flexibility. Strikingly, Neu Aralar-KO mice displayed a clear impairment in sociability, which together with the observed hyperactivity, identifies neuronal Aralar deficiency as a determinant of behavioral abnormalities relevant to ASD (Silverman et al., 2010; Kwon et al., 2022; Giliberti et al., 2025).

### 3. Neuronal Aralar ablation remodels dendritic spine morphology and striatal transcriptomics

Alterations in synaptic function, and changes in dendritic spine density and morphology have been closely linked to neurodevelopmental and neuropsychiatric disorders characterized by impaired social behavior, including ASD in patients and mouse models (Zoghbi et al., 2003; Hutsler and Zhang, 2010; Penzes et al., 2011; Phillips and Pozzo-Miller, 2015; Akhgari et al., 2024; Reyes-Lizaola et al., 2024). Given the critical role of the striatum in regulating social behavior (Yin and Knowlton, 2006; Báez-Mendoza and Schultz, 2013), and the sociability deficits observed in Neu Aralar-KO mice, we focused our analyses on this brain region. Specifically, we investigated whether neuronal Aralar deficiency affects dendritic spine density and morphology in 7-8-month-old Neu Aralar-KO mice, an age at which social deficits were apparent (Figure 2O). To this end, brain slices were labeled with the lipophilic fluorescent dye DiI. Ǫuantification of dendritic spine density revealed no differences between Neu Aralar-KO and control mice (Figure 3A-B); but morphometric analyses showed significant alterations in dendritic spine structure (Figure 3C-F). Spine head surface area and neck length in striatum were increased by 12% and 35%, respectively in Neu Aralar-KO striata. These changes resulted in a significantly reduced head surface-to-neck length ratio (Figure 3E), indicative of larger spine heads (Figure 3F) positioned closer to the synaptic cleft. Together, these morphological changes in striatal dendritic spines may be indicative of strengthened synapsis with larger synaptic surface and better voltage compartmentalization (Chidambaram et al., 2019). Morphological alterations in dendritic spines were also observed in cortical neurons from Neu Aralar-KO mice; with no changes in density (Figure S2A-F).

**Figure 3.**
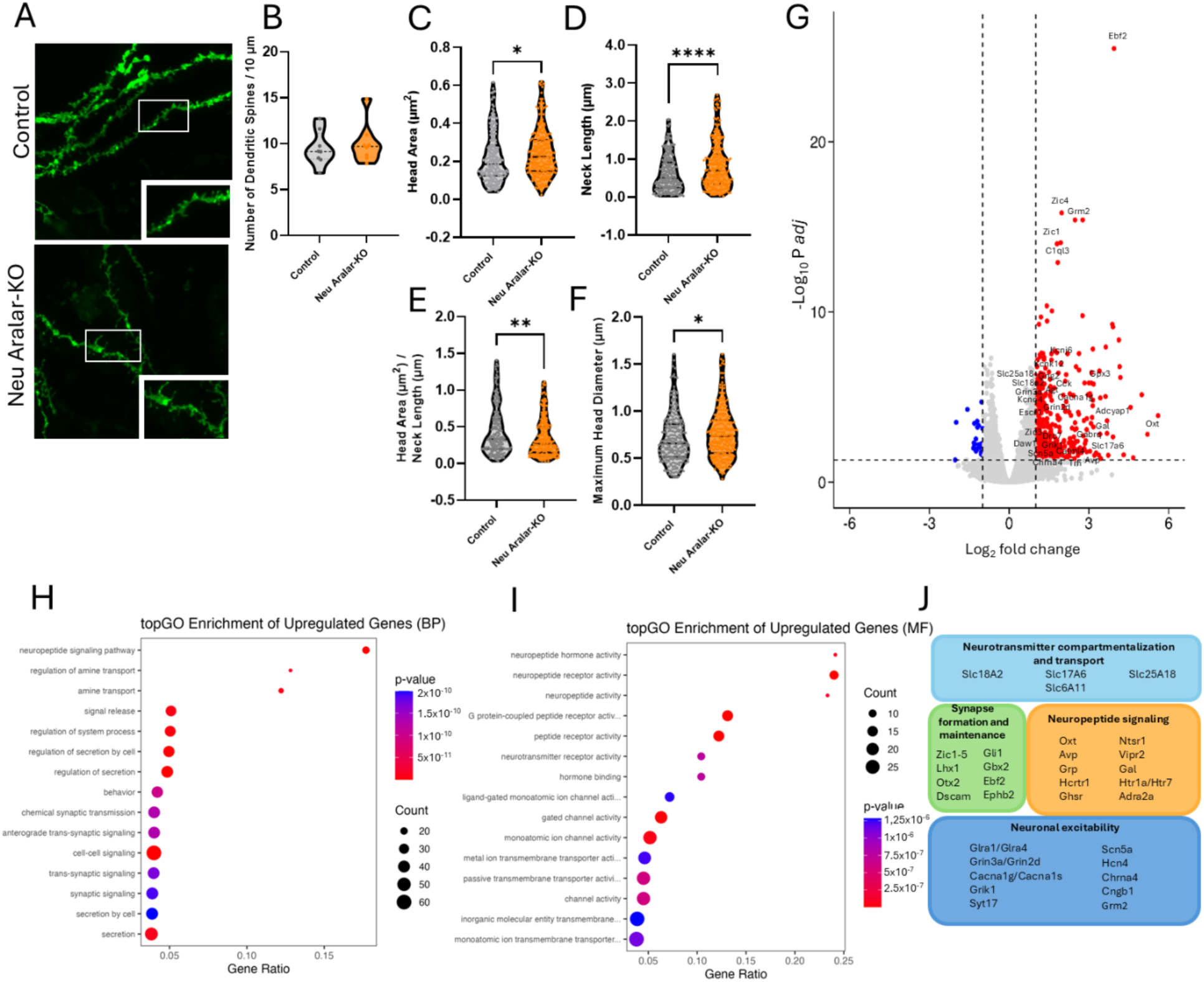
Aralar ablation remodels dendritic spine morphology and dysregulates striatal neurotransmission-related transcriptional programs. (**A**) Representative images of dendritic spines in striatal neurons from Control (top) and Neu Aralar-KO (bottom) mice following DiI microcrystal labelling; insets show the boxed regions at higher magnification. Ǫuantification of (**B**) dendritic spine density, (**C**) head area, (**D**) neck length, (**E**) head area-to-neck length ratio and (**F**) maximum spine head diameter. (**G**) Volcano plot showing differentially expressed genes (DEGs) between the striata of Control and Neu Aralar-KO mice. Blue dots indicate significantly downregulated genes (adjusted p < 0.05, log_2_FC < −1), red dots significantly upregulated genes (adjusted p < 0.05, log_2_FC > 1) and grey dots non-significant genes; selected genes are labelled. Gene Ontology (GO) enrichment analysis of upregulated DEGs (top 15 terms) categorised by (**H**) biological process (BP) and (**I**) molecular function (MF); dot size indicates the number of genes in the term and dot colour the enrichment p value. Enrichment significance was calculated using Fisher’s exact test (topGO). (**J**) Upregulated genes in the Neu Aralar-KO striatum grouped into functional modules: neurotransmitter compartmentalisation and transport, synapse formation and maintenance, neuropeptide signalling, and neuronal excitability. Data in B–F are presented as mean ± SEM. Statistical analysis was performed using unpaired Student’s *t*-test (*p < 0.05;\*\***p < 0.01**; ****p < 0.0001). n = 3 mice per genotype.

To obtain an unbiased overview of the molecular effects of Aralar ablation in the ventral striatum, we performed a transcriptomic analysis in Neu Aralar-KO mice and their control littermates. We analyzed differentially expressed genes (DEGs) in both genotypes (Figure 3G-J). Genes were considered differentially expressed if they met the criteria of adjusted p-value < 0.05 and absolute log_2_ fold change (|log_2_Fold|) > 1, acknowledging that not all cells in the striatal region were recombinant. The analysis identified a total of 433 DEGs in the striatum, of which 409 were significantly upregulated in Neu Aralar-KO mice compared to controls (Figure 3G). Gene Ontology (GO) enrichment analysis revealed that the most significantly upregulated biological processes are related to neuropeptide signaling, synaptic transmission and signaling, secretion and behavior (Figure 3H). Accordingly, GO enrichment analysis of upregulated genes highlighted significant overrepresentation of molecular functions related to neuropeptide hormone activity, neuropeptide receptor activity, G protein-coupled peptide receptor activity, peptide/neurotransmitter receptor activity, and gated ion channel activity (Figure 3I). Additional enriched categories included transmembrane transporter activity and other ion channel-related functions, indicating broad effects on neuronal signaling and membrane excitability.

Upregulated genes could be organized into distinct functional modules associated with: 1) “Neurotransmitter compartmentalization and transport”, 2) “Synapse formation and maintenance”, 3) “Neuronal excitability”, and 4) “Neuropeptide signaling” (Figure 3J). Of particular relevance to the module of “Neurotransmission compartmentalization and transport”, are genes such as *slc18a2* and *slc17aC* encoding the vesicular monoamine and glutamate transporters 2 (VMAT2 and VGLUT2), respectively; *slc25a18*, coding for the mitochondrial glutamate carrier 2 (GC2) which, as Aralar/*slc25a12,* transports glutamate into mitochondria; and, *slcCa11*, encoding a transporter involved in the uptake of extracellular GABA in astrocytes. A prominent module for “Synapse formation and maintenance” includes the transcription factor *Ebf2* (Martínez-Estrada et al., 2026); *Ephb2* (Ethell et al., 2001), or *Dscam* (Down Syndrome Cell Adhesion Molecule) (Li et al., 2009; Maynard and Stein, 2012).

Other DEGs can be related to Neuronal excitability, including among others: *Grin3a*, *Grin2d*, and *Grm2*, encoding NMDA receptor subunits and a metabotropic Glu receptor, respectively; *Grik1*, a KA receptor subunit and *Syt17*, a synaptotagmin that regulates postsynaptic AMPA receptor density and contributes to synaptic plasticity (Ruhl et al., 2019); as well as several ion channels (*Cacna1, Scn5a, Hcn4*). Interestingly, genes in neuropeptide signaling (Figure 3H-J) were especially present within the upregulated DEGs. The coordinated upregulation of *Ntsr1*, *Hcrtr1*, *Htr1a* and *Htr7*, encoding the neurotensin, hypocretin/orexin type 1, and serotonin 1A and 7 receptors respectively, suggests an adaptive remodelling of neuromodulatory systems involved in social motivation and arousal (Sakurai, 2007; Hedlund, 2009; Tan et al., 2020; Singhal et al., 2023). Notably, oxytocin and serotonin act in concert within the nucleus accumbens to support social reward (Dölen et al., 2013), a circuit in which neurotensin signalling also modulates dopamine release (Singhal et al., 2023).

### 4. Neuronal Aralar deficiency causes alteration in DA metabolism and increased oxidative stress in striatum

The impaired sociability in Neu Aralar-KO mice (Figure 2O) could potentially be associated with alterations in excitatory/inhibitory neurotransmitter balance (Weiss et al., 2021) and/or DAergic signaling within the mesocorticolimbic circuitry involved in social reward and motivation (Figure 4H; Gunaydin et al., 2014). To explore these possibilities, we quantified amino acid and monoamine levels in ventral striatal and cortical regions from 7–8 month-old mice (Figure 4 A-H and Figure S2G-K, respectively). Interestingly, in contrast to global Aralar KO mice (Llorente-Folch et al., 2013b; Peréz-Liébana et al., 2020) neither Glu nor GABA levels were significantly altered in the striatum or cortex of Neu Aralar-KO mice (Figure 4B,4C and Figure S2G, respectively). In the cortex of Neu Aralar-KO mice, there was a decrease in Asp and Serine (Ser) levels (Figure S2H, S2I) as in the cortex of global Aralar-KO mice (Pardo et al., 2011), while monoamine levels were unchanged (Figure S2G). In the Neu Aralar-KO striatum, Asp and Ser levels were also reduced, but only Ser decrease reached statistical significance (33% decrease; Figure 4D, 4E). Serotonin (5-HT) and its byproduct (5-hydroxyindoleacetic acid; 5-HIAA) showed no changes (not shown). However, DA levels were significantly reduced in Neu Aralar-KO striata (38% with respect to control); and the DOPAC (3,4-dihydroxyphenylacetic acid; a DA degradation product) to DA ratio increased by ≈ 3-fold, indicating enhanced intraneuronal DA metabolism (Figure 4F,4G), as observed in the global Aralar KO mouse (Llorente-Folch et al., 2013b). Intriguingly, CaMKIIα-Cre-mediated recombination was largely restricted to cortical and striatal neurons, (Figure 1D-F), but mesencephalic DAergic neurons, identified by tyrosine hydroxylase (TH) immunoreactivity, exhibited only very limited recombination (aprox. 15% of TH-positive neurons co-express the ZsGreen reporter). This was particularly evident in the VTA, a key DAergic mesencephalic region involved in social reward processing, where recombination was not detected (Figure 4H). These observations indicate that *Aralar* expression is largely preserved in midbrain DAergic neurons and, consequently, that the disruption of DAergic function in ventral striatal terminals from VTA neurons takes place despite its presence in these neurons. Therefore, this is likely due to non-cell autonomous effects.

**Figure 4.**
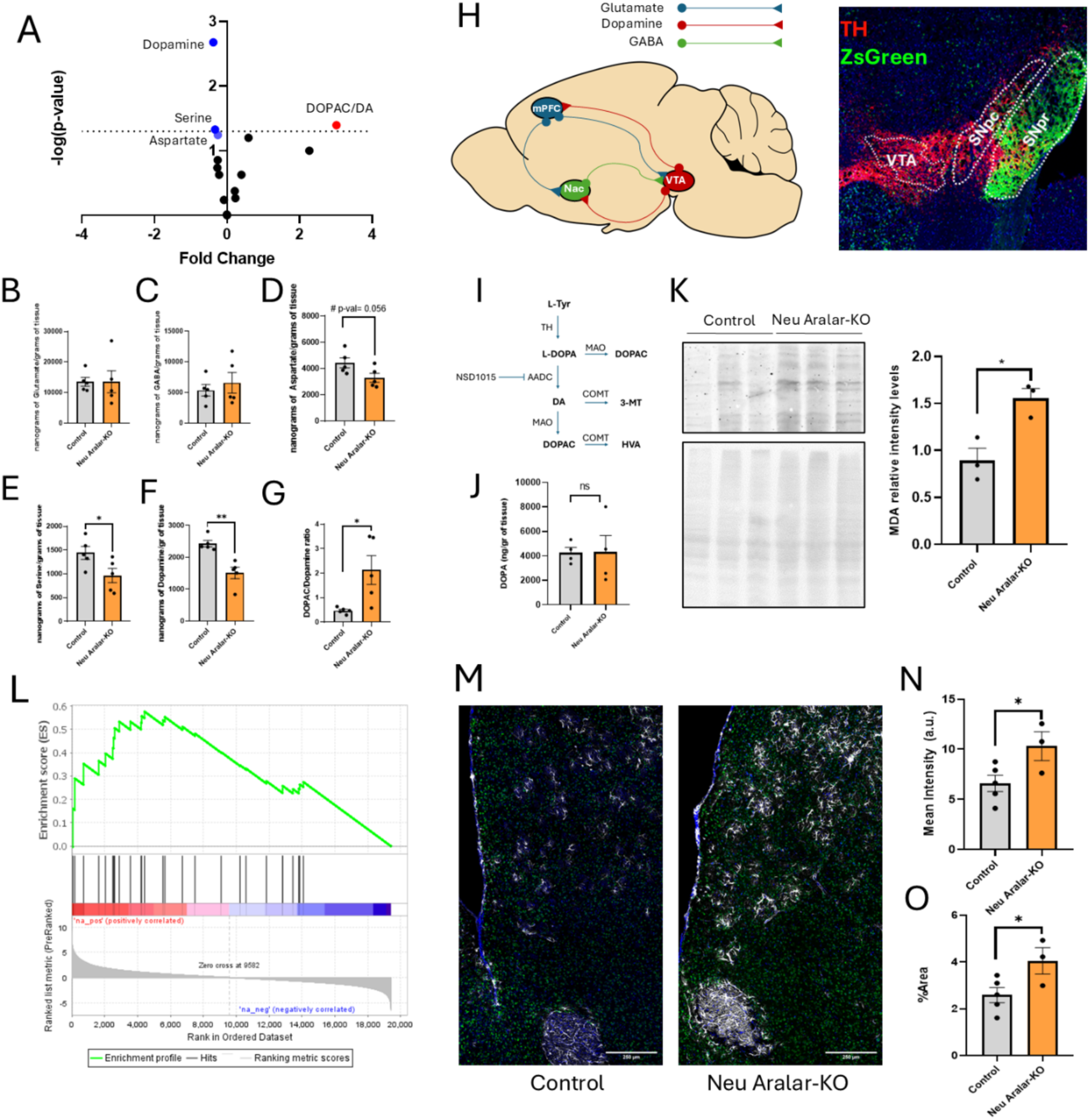
Aralar ablation promotes oxidative stress-associated dopamine mishandling in the striatum. (**A**) Volcano plot of amino acid and monoamine alterations in the striatum of Neu Aralar-KO relative to Control mice, plotted as −log(p value) against fold change. Metabolites significantly decreased and increased in the Neu Aralar-KO are shown in blue and red respectively and are labelled; the dotted horizontal line indicates the significance threshold (p = 0.05). Striatal concentrations of (**B**) glutamate, (**C**) GABA, (**D**) aspartate, (**E**) serine and (**F**) dopamine, expressed as nanograms per gram of tissue, and (**G**) the DOPAC/dopamine ratio. (**H**) Schematic representation of the dopaminergic and corticostriatal pathways analysed (mPFC, medial prefrontal cortex; NAc, nucleus accumbens; VTA, ventral tegmental area), together with a representative image of TH immunostaining (red) and the ZsGreen recombination reporter (green) in the ventral midbrain, showing that VTA and SNpc dopaminergic neurons do not undergo recombination. (**I**) Schematic overview of dopamine synthesis from L-tyrosine (L-Tyr) and of the experimental design for AADC inhibition: mice received NSD1015 (100 mg/kg, i.p.) and striatal tissue was collected 60 min later. L-DOPA, L-3,4-dihydroxyphenylalanine; TH, tyrosine hydroxylase; AADC, aromatic L-amino acid decarboxylase; DA, dopamine; MAO, monoamine oxidase; COMT, catechol-O-methyltransferase; DOPAC, 3,4-dihydroxyphenylacetic acid; 3-MT, 3-methoxytyramine; HVA, homovanillic acid. (**J**) Striatal DOPA accumulation following NSD1015 administration, used as a read-out of *in vivo* TH activity. (**K**) Representative immunoblot of malondialdehyde (MDA) protein adducts in Control and Neu Aralar-KO striata, with the corresponding total-protein loading control, and densitometric quantification. (**L**) Gene set enrichment analysis of the KEGG glutathione metabolism pathway, showing the normalised enrichment score (NES) and FDR q value. (**M**) Representative GFAP immunostaining in the ventral striatum of Control (left) and Neu Aralar-KO (right) mice. Ǫuantification of (**N**) GFAP mean fluorescence intensity and (**O**) GFAP-positive area. Data are presented as mean ± SEM (n = 5 mice for Control and n = 5 mice Neu Aralar-KO for B–G; n = 4 mice per genotype for J; n = 3 per genotype for K; n = 5 Control and n = 3 Neu Aralar-KO for M–O). Statistical analysis was performed using unpaired Student’s *t*-test; n.s., not significant; # denotes a non-significant trend (p = 0.056); *p < 0.05, **p < 0.01. Scale bars, 250 µm (M).

Low DA levels in Aralar-deficient striatum point to increased intraneuronal DA degradation. However, the reduction in DA levels (Figure 4B) could also result from impaired synthesis by TH in DAergic terminals. To assess TH activity *in vivo* (Caudle et al., 2007), aromatic L-amino acid decarboxylase (AADC), which converts DOPA into DA (Figure 4I), was pharmacologically inhibited, and the accumulation of striatal DOPA used as a readout of TH activity. No differences in DOPA accumulation were detected between genotypes (Figure 4J), indicating that the reduction in DA levels is not attributable to an impaired DA synthesis. Mishandling of striatal DA, reflected by the elevated DOPAC/DA ratio, in Neu Aralar-KO mice (Figure 4G), indicates enhanced intracellular DA catabolism via monoamine oxidase (MAO), a process that generates H₂O₂ as a by-product (Spina and Cohen, 1989). This process is expected to generate oxidative stress within the striatum, a region in which mesencephalic DAergic terminals are particularly vulnerable to such insults (Burbulla et al., 2017; Surmeier et al., 2017). Global Aralar-KO mice have a similar defect in DA handling which was attributed to oxidative stress in the DAergic neuron (Llorente-Folch et al., 2013b).

To determine whether neuronal Aralar deficiency induces oxidative stress, we quantified MDA-modified proteins by Western blot, a well-established marker of lipid peroxidation and oxidative damage (Giera et al., 2012; Dharmajaya et al., 2022). MDA levels increased by 66% in the striatum of Neu Aralar-KO mice compared with controls (Figure 4K). Consistently, Gene Set Enrichment Analysis (GSEA) revealed significant enrichment of glutathione metabolism pathways in the Neu Aralar-KO striatum (Figure 4L). Several genes involved in cellular protection against oxidative stress were also strongly upregulated, including Aldh3b2, which detoxifies reactive aldehydes generated during lipid peroxidation and has been reported to colocalize with astrocytic markers (Han et al., 2022); and Gpx3, a secreted extracellular glutathione peroxidase, both showing approximately 8-fold increases in expression. The induction of these antioxidant pathways was accompanied by a marked increase in GFAP immunoreactivity (Figure 4M-O), consistent with the well-established astrocytic response to neuron-derived oxidative stress (Fulton et al., 2021; Lee et al., 2026). Notably, the robust upregulation of Gpx3 suggests elevated levels of diffusible extracellular ROS. Together, these findings indicate that neuronal Aralar deficiency triggers oxidative stress that extends beyond neurons and impacts neighboring astrocytes. Given the high intrinsic vulnerability of DAergic terminals to oxidative damage, this altered redox environment is likely to contribute to the enhanced oxidative catabolism of DA observed in the striatum.

### 5. β-Hydroxybutyrate (BHB) restores sociability, metabolic alterations and oxidative stress in neuronal Aralar/AGC1 deficiency: BHB-mediated rescue mechanisms

After demonstrating the protective role of ketogenic diet (KD) in patients with AGC1-deficiency (Dahlin et al., 2015; Pfeiffer et al., 2020); and that of BHB in global Aralar-KO mice (Pérez-Liébana et al., 2020), we interrogated whether *in vivo* BHB treatment could reverse the impairments observed in Neu Aralar-KO mice. BHB may exert its protective effects in brain through several mechanisms, a) requiring BHB metabolism: (1) restoring mitochondrial energy metabolism; (2) enhancing cytosolic Asp synthesis in Aralar-KO neurons; (3) reducing ROS levels; or, b) independent of BHB metabolism: (4) via epigenetic regulation through histone deacetylase inhibition (Shimazu et al., 2013), and/or β-hydroxybutyrylation in the brain (Cornuti et al., 2023); or 5) via BHB receptors in glia (Rahman et al., 2014).

To study the effects of BHB, Neu Aralar-KO and control mice were injected daily with i.p. BHB (520 mg/Kg; 5 mM final concentration), to mimic the BHB concentration found in the blood of AGC1 deficient patients treated with KD (Dahlin et al., 2015). Social behavior was assessed prior to and after 15 days of BHB administration (Figure 5A, B). To prevent recognition bias, a new set of unfamiliar stimulus mice was used during the second test after BHB treatment (Figure 5B). At baseline, Neu Aralar-KO mice displayed a significantly reduced social DI of 0.52 indicating no preference between a social stimulus (novel mouse) and an inanimate object; whereas control mice showed a typical social preference (ID, 0.64). Remarkably, after BHB treatment, the social DI in Neu Aralar-KO mice significantly increased to 0.59, abolishing genotype differences observed in untreated animals (Figure 5B and 2O).

**Figure 5.**
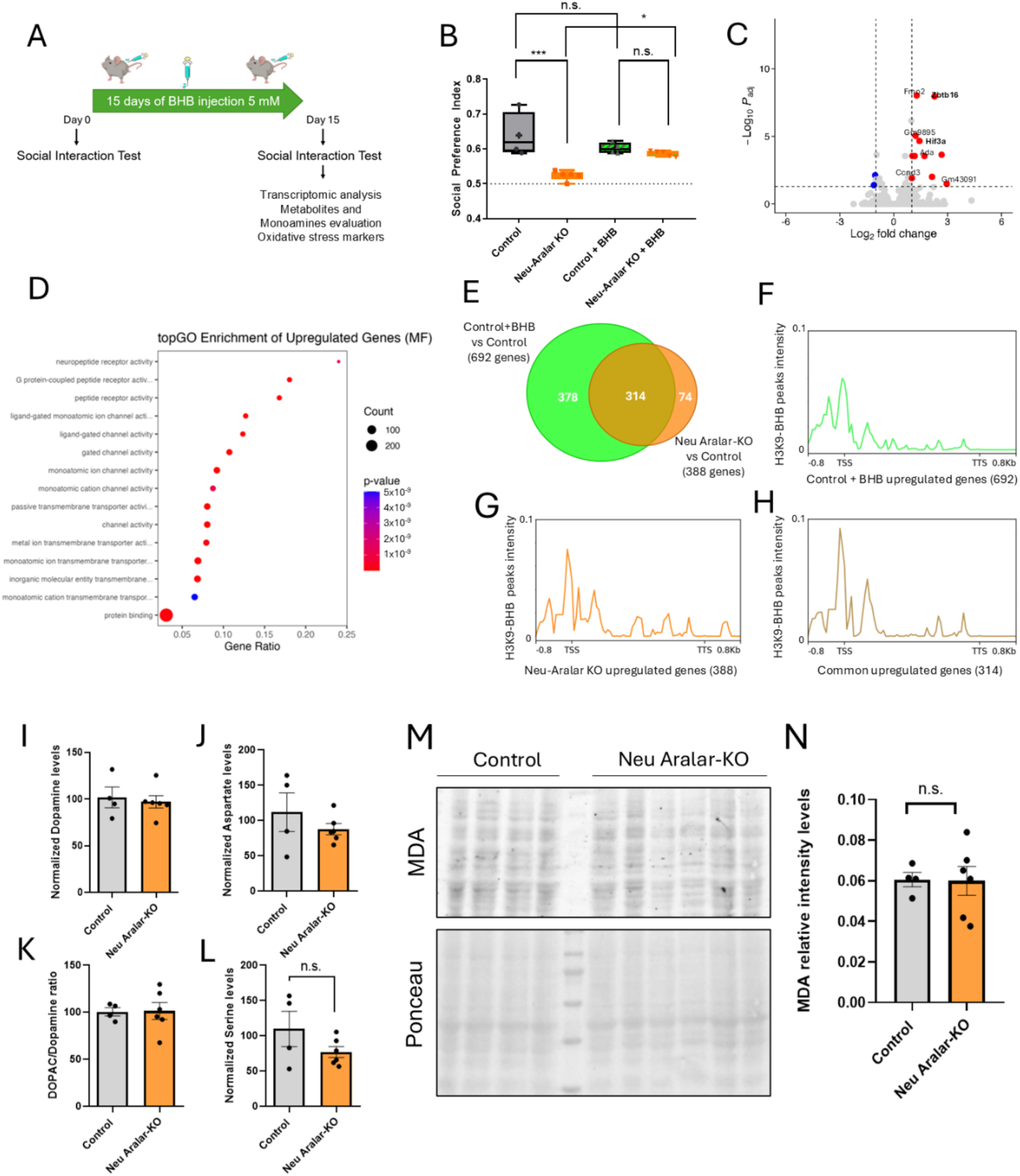
β-Hydroxybutyrate treatment rescues social behaviour and restores metabolic and redox homeostasis. (**A**) Experimental design for β-hydroxybutyrate (BHB) treatment: mice received daily BHB injections (5 mM) for 15 days; social behaviour was assessed before (day 0) and after (day 15) treatment, after which striatal tissue was collected for transcriptomic analysis, quantification of metabolites and monoamines, and measurement of oxidative stress markers. (**B**) Social preference index in the three-chamber social interaction test for Control, Neu Aralar-KO, Control + BHB and Neu Aralar-KO + BHB groups (Control, n = 4; Neu Aralar-KO, n = 6). Statistical analysis was performed using repeated measurement ANOVA with *post hoc* correction for multiple comparisons. (**C**) Volcano plot showing differentially expressed genes (DEGs) in BHB-treated Neu Aralar-KO mice compared with untreated Neu Aralar-KO mice (adjusted p < 0.05, |log_2_FC| > 1). (**D**) Top 15 enriched Gene Ontology (GO) terms in the molecular function (MF) category for genes upregulated in BHB-treated Control mice relative to untreated Control mice. (**E**) Venn diagram showing the overlap between the genes upregulated in the Control + BHB *versus* Control comparison (692 genes) and those upregulated in the Neu Aralar-KO *versus* Control comparison (388 genes). (**F–H**) Metaplots of H3K9-bhb ChIP-seq signal intensity, taken from publicly available data (Cornuti et al., 2023), over the gene sets defined in (E): genes upregulated in Control + BHB *versus* Control (**F**), genes upregulated in Neu Aralar-KO *versus* Control (**G**) and genes common to both comparisons (**H**). Signal is displayed from 0.8 kb upstream of the transcription start site (TSS) to 0.8 kb downstream of the transcription termination site (TTS). The ChIP-seq dataset represents H3K9-bhb peaks upregulated under fasting relative to *ad libitum* feeding in mouse cortex. HPLC-based quantification of (**I**) dopamine, (**J**) aspartate, (**K**) the DOPAC/dopamine ratio and (**L**) serine in BHB-treated Control and Neu Aralar-KO mice. The same animals were used for behavioral and biochemical analyses (Control, n = 4; Neu Aralar-KO, n = 6). (**M**) A representative immunoblot of malondialdehyde (MDA) protein adducts in striata of BHB-treated Control and Neu Aralar-KO mice; Ponceau staining was used as loading control. (**N**) Densitometric quantification of MDA immunoreactivity normalised to the corresponding loading control. Statistical analyses for panels I–N were performed using unpaired Student’s *t*-test. Data are presented as mean ± SEM. n.s., not significant; *p < 0.05; ***p < 0.001.

To assess the impact of BHB on striatal transcriptional changes in Neu Aralar-KO mice, DEG analysis was performed (Figure 5C). BHB effect on transcriptomics of Neu Aralar-KO mice was minimal, with only 13 DEGs identified (11 upregulated). The low number of DEGs precluded robust GO enrichment analysis. Among the few altered genes in Neu Aralar-KO striata are those encoding the transcription factors Hif3α and Zbtb16, both also upregulated by BHB in control mice. Notably, Zbtb16 KO mice have impaired social behavior, and Zbtb16 downregulation has been found in ASD patients (Usui et al., 2021), providing a link to ASD-related social and cognitive behaviors. The limited transcriptional response to BHB treatment in Neu Aralar-KO mice was unexpected, since it is well-known the transcriptional effect of BHB in control cells (Shimazu et al., 2013; Cornuti et al., 2023). Also, it was surprising that BHB in control striata, upregulated genes, belonging to enriched GO terms for molecular functions, very similar to those significantly upregulated in striata from Neu Aralar-KO mice (11/15 terms were common; compare Figure 5D to 3I).

To investigate this unexpected result, we compared the DEG sets identified in control striata following BHB treatment and in untreated Neu Aralar-KO striata (Figure 5E). Strikingly, nearly half of the genes that were upregulated following BHB treatment in control striata were already upregulated in the untreated Neu Aralar-KO striatum (313 of 690 genes), suggesting that Aralar deficiency may partially recapitulate a transcriptional program associated with BHB exposure.

To further explore this overlap, we compared the DEGs from this work to publicly available C57BL/6J mice cortex H3K9-bhb ChIP-seq profiles from Cornuti et al. (2023). H3K9-bhb refers to β-hydroxybutyrylation of histone H3 at lysine 9, a histone post-translational modification associated with activation of gene expression (Xie et al., 2016). The published dataset included genomic regions showing increased H3K9-bhb signal under fasting conditions compared with ad libitum feeding in mice cortex. As fasting is known to increase ketogenesis, its effects on H3K9 may mimic those of BHB. Metaplot analysis revealed enrichment of H3K9-bhb signal around the TSSs (Transcription starting sites) of genes upregulated following BHB treatment in control mice (Figure 5F) and in Neu Aralar-KO upregulated genes (Figure 5G). Notably, common genes between the two DEG sets exhibited specific H3K9-bhb enrichment around TSS regions (Figure 5H). Control pileup plots with shuffled regions showed no signal enrichment around the TSS (Figure S3). Together, these findings reveal that a great portion (1/2) of genes differentially expressed/activated upon BHB treatment in control striata are also differentially expressed in Neu Aralar-KO. This convergence suggests that neuronal Aralar deficiency may engage transcriptional programs that partially overlap with those associated with BHB exposure and fasting-induced H3K9-bhb enrichment.

The recovery of social behavior deficits following BHB treatment (Figure 5B) prompted us to investigate whether this was accompanied by a correction of the metabolic alterations in the Neu Aralar-KO striata. While short-term BHB treatment (5 days) in global Aralar-KO mice did not fully normalize metabolic defects (Perez-Liébana et al., 2020), the current study employed a less severely affected mouse model with a longer treatment period (15 days) and a higher BHB dose comparable to that reported in an AGC1-deficient patient following an effective KD (Dahlin et al., 2015). We measured amino acids and monoamines in striatal biopsies in BHB-treated Neu Aralar-KO and control mice (9–10 months old). Strikingly, after 15 days of BHB administration, the metabolic abnormalities were no longer present in Neu Aralar-KO striata (Figure 5I-L). Given the potential association between the metabolic defects and increases in oxidative stress, we wondered whether BHB was able to recover also the increase in MDA levels in Neu Aralar-KO striatum (Figure 4K). In fact, MDA immunostaining performed on the BHB-treated Neu Aralar-KO and control striata revealed no differences between both experimental groups (Figure 5M, N). These results suggest a reduction of oxidative stress upon BHB treatment, which aligns with the recovery in metabolites, and is also consistent with our previous results (Pérez-Liébana et al, 2020).

Altogether, our results support a model in which Aralar deficiency in NAc MSNs causes a metabolic impairment that increased ROS production, leading through non-cell-autonomous mechanisms to DAergic dysfunction in VTA terminals, glial activation, and impaired sociability in NeAralar-KO mice (Figure 6). The ability of BHB to rescue oxidative stress, DAergic and behavioral alterations, despite its limited transcriptional impact in Neu Aralar-KO striatum, suggests that restoration of metabolic homeostasis and redox balance is the primary mechanism underlying its therapeutic action.

**Figure 6.**
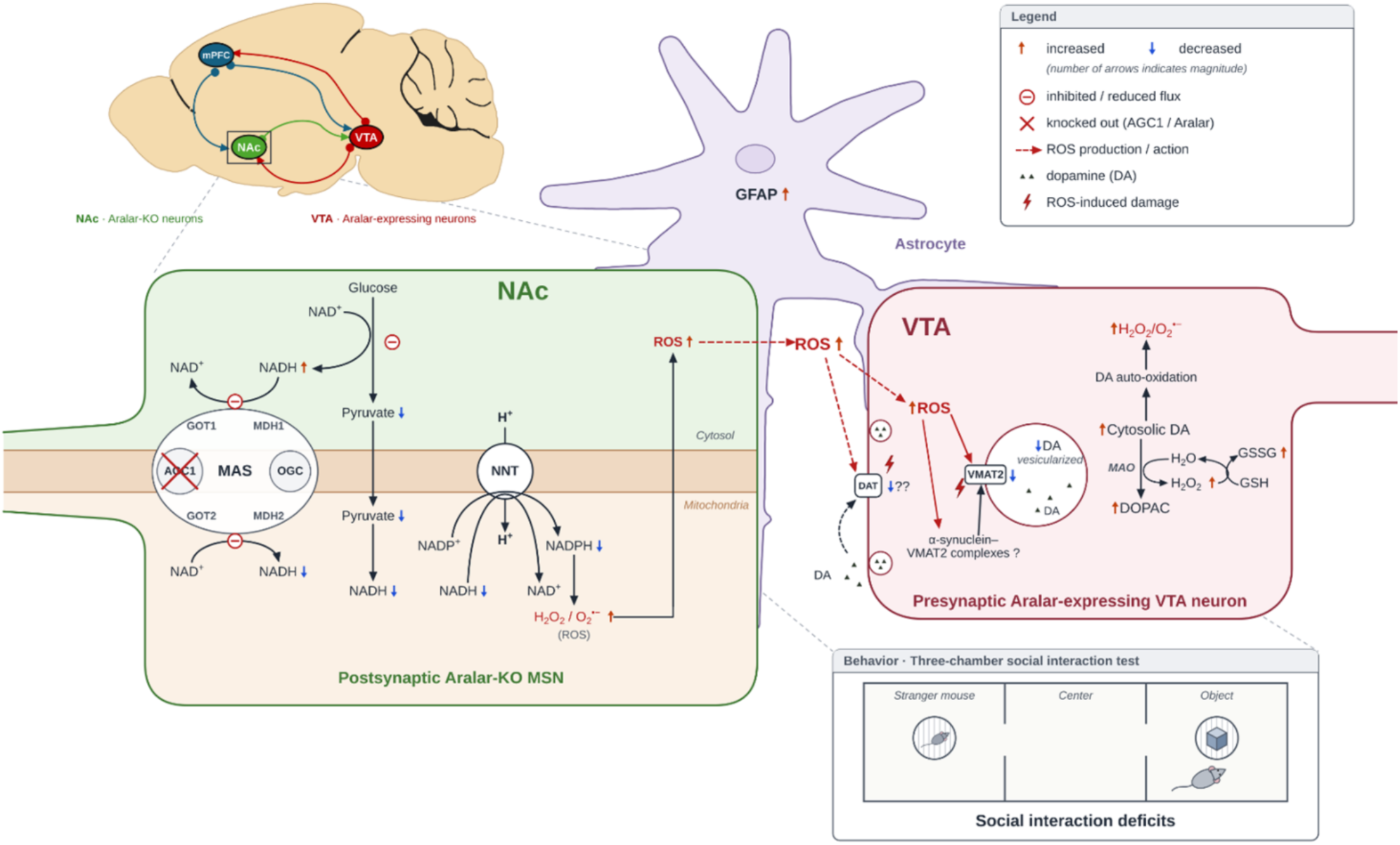
Proposed mechanism linking neuronal Aralar/AGC1 deficiency to redox imbalance, dopaminergic dysfunction and impaired sociability. Schematic representation of the mesocorticolimbic pathway, indicating Aralar-deficient neurons in the nucleus accumbens (NAc) and Aralar-expressing dopaminergic neurons in the ventral tegmental area (VTA). AGC1 deficiency in NAc medium spiny neurons (MSNs) reduces glycolytic flux and pyruvate production. The consequent decrease in mitochondrial pyruvate availability and in MAS activity lowers mitochondrial NADH levels, so that NNT-dependent NADPH generation is reduced and ROS detoxification capacity is compromised. ROS generated in Aralar-KO MSNs may diffuse into the extracellular space and affect neighbouring cells, including astrocytes, which show increased GFAP immunoreactivity, and mesencephalic dopaminergic terminals arising from the VTA. In these dopaminergic neurons, ROS may directly damage the DAT and VMAT2 transporters or promote the formation of VMAT2–α-synuclein complexes, reducing the vesicular dopamine pool. The resulting accumulation of free cytosolic dopamine is more susceptible to auto-oxidation and to MAO-mediated metabolism, both of which generate further ROS. Metabolic impairment of Aralar-deficient NAc MSNs therefore induces adaptive changes in dopamine handling in Aralar-expressing VTA neurons, revealing a non-cell-autonomous effect, together with enhanced astrogliosis. Altered dopaminergic signalling within the mesolimbic reward circuitry ultimately compromises social reward processing, resulting in the reduced sociability observed in Neu Aralar-KO mice in the three-chamber social interaction test. Arrows indicate increased (↑) or decreased (↓) levels; ⊖ indicates inhibited or reduced flux and ✕ indicates the knocked-out carrier (Aralar/AGC1).

## DISCUSSION

The present study provides new insights into the molecular and cell-specific functions of Aralar, a mitochondrial carrier known to play a pivotal role in neuronal metabolism, including mitochondrial respiration, lactate utilization, and Asp synthesis; and glial functions as Gln synthesis or postnatal myelination (Pardo et al., 2022). While the essential contribution of Aralar to brain function has been established by the severe neurological phenotype observed in AGC1-deficient patients and global KO mouse models (Wibom et al., 2009; Jalil et al., 2005; Llorente-Foch et al., 2013b), the phenotypic complexity and widespread Aralar expression has made it difficult to define the mechanisms and cell types driving the diverse manifestations of Aralar deficiency. Our findings help address this gap by focusing on neurons, the brain cell type expressing the highest levels of Aralar and likely the most dependent on its function. To do this, we generated and characterized a neuronal-specific Aralar-KO mouse model. Intriguingly, unlike global Aralar-KO mice, which die at PND20 (Jalil et al., 2005), selective ablation of Aralar in mature neurons had no detectable effect on somatic growth or survival. Remarkably, however, these mice displayed selective behavioral and neurochemical alterations, indicating that neuronal Aralar deficiency disrupts discrete aspects of brain function.

### 1. Neuronal Aralar deficiency impairs sociability but preserves memory

Behavioral assessment of Neu Aralar-KO mice revealed no detectable deficits in memory or cognitive flexibility, despite the established role of lactate as an energetic fuel for memory formation (Suzuki et al., 2011; Magistretti and Allaman, 2018) and the impaired ability of Aralar-KO neurons to use lactate efficiently (Pardo et al., 2006; Llorente-Folch et al., 2016). Interestingly, Neu Aralar-KO mice exhibited marked hyperactivity and reduced sociability. The latter findings are particularly noteworthy, as it is consistent with the proposed involvement of *SLC25A12/Aralar* in ASD susceptibility (Ramoz et al., 2004; Lepagnol-Bestel et al., 2008, Durdiaková et al., 2014, Aoki et al., 2016, Yao et al., 2021; Ghavi et al., 2024).

We hypothesized that the prevention of lactate oxidation by Aralar deficiency in neurons could affect memory acquisition given the demonstrated role of lactate in memory formation (Suzuki et al., 2011; Magistretti and Allaman, 2018). However, memory functions in Neu Aralar-KO mice were found to be intact across diverse behavioral assays (Figure 2). The ANLS hypothesis for synaptic plasticity and memory, posits that lactate produced by astrocytes in response to neuronal activity, is used primarily as a neuronal metabolic fuel with signaling properties. However, recent findings question the role of lactate as neuronal metabolic fuel in the presence of glucose (Söder et al., 2025; Rauseo et al., 2026), suggesting that the relevant role of lactate within ANLS is probably related with signaling and/or posttranslational mechanisms rather than metabolic. As Aralar-KO neurons are impaired for lactate metabolic utilization (Pardo et al., 2006) but not signaling, this may explain memory preservation in Neu Aralar KO mice.

Notably, impaired sociability emerged as the hallmark behavioral phenotype of Neu Aralar-KO mice. This is important given the relationship between Aralar and ASDs, based on associations between SNPs in *Slc25a12* gene in ASD (Ramoz et al., 2004; Lepagnol-Bestel et al., 2008; Durdiaková et al., 2014; Aoki et al., 2016; Yao et al., 2021) and with decreased levels of Aralar in brains of ASD individuals (Ghavi et al., 2024). Of note, the three-chamber social interaction test used in this study was conducted in female mice—a group traditionally underrepresented in ASD research— highlighting the need to consider sex differences in neurodevelopmental studies (Bhargava and Ashwin, 2025).

Since the NAc functions as a central integrative hub for social reward, under the modulatory influence of VTA-derived DAergic inputs (Grace et al., 2007; Gunaydin et al., 2014), we performed further characterization of the Neu Aralar-KO ventral striata. Notably, DA was the only neurotransmitter significantly altered in these mice. In contrast to global Aralar-KO animals, which displayed also marked reductions in both Glu and GABA levels (Llorente-Folch et al., 2013b). Nevertheless, transcriptomic profiling in Neu Aralar-KO striata revealed dysregulation of genes associated with glutamatergic and GABAergic neurotransmission, suggesting functional remodeling of these networks despite preserved steady-state neurotransmitters levels. DAergic homeostasis, however, was profoundly disrupted in Neu Aralar-KO striatum, as evidenced by reduced DA levels and a marked increase in DA turnover (elevated DOPAC/DA ratios). Besides, DAergic dysfunctions were previously reported in the global Aralar-KO mice (Llorente-Folch et al 2013b; Perez-Liebana et al., 2020). Several non-mutually exclusive mechanisms could account for the reduction in DA levels, including: (i) impaired DA synthesis resulting from oxidative damage to TH; (ii) enhanced intraneuronal DA degradation through MAO-dependent metabolism and/or auto-oxidation; and (iii) increased utilization of DA as a substrate for MAO-driven mitochondrial ETC activity to support neuronal bioenergetics (Graves et al., 2020). Our data argue against impaired DA synthesis as the primary mechanism, as neither TH levels nor activity differed between genotypes. Instead, Neu Aralar-KO mice exhibited clear evidence of increased intraneuronal DA metabolism accompanied by elevated oxidative stress. Collectively, these findings indicate that neuronal Aralar deficiency promotes the accumulation of a larger cytosolic pool of free DA that is particularly susceptible to MAO-dependent degradation and auto-oxidation, thereby reducing tissue DA without altering its rate of synthesis.

### 2. Impaired function of DA neurons is non-cell-autonomous

Our study reveals a surprising finding. That the removal of Aralar in MSN of the ventral striatum results in a dramatic disruption of DA handling in neighbouring DAergic terminals from VTA neurons in which Aralar expression is maintained. In other words, that the defect in the DA handling in VTA terminals is not cell-autonomous. The most likely explanation of this finding is based on the observation that the lack of Aralar in brain neurons results in an increase in oxidative stress response when exposed to excitatory stimuli (Llorente-Folch et al., 2016) and a drop in the GSH/GSSG ratio in brain of Aralar-KO mice (Llorente-Folch et al., 2013b).

At the neuronal level, glutamate exposure to cortical neurons induced an early appearance of 8-OHdG, an oxidized nucleoside formed after oxidative stress, in nuclear and especially mitochondrial DNA which was blocked in the presence of lactate in wild type but not in MAS-defective aralar-KO neurons (Llorente-Folch et al., 2016). It is unlikely that the increased oxidative stress response in Aralar-KO neurons is due to an intrinsic overproduction of superoxide (O_2_^-^) by their electron transport chain, but rather due to an impairment in their mitochondrial antioxidant systems. Indeed, NADH availability in mitochondria is required to form NADPH via nicotinamide nucleotide transhydrogenase (NNT) which catalyses the reduction of NADP+ at the expense of NADH oxidation coupled to inward proton translocation across the inner mitochondrial membrane. NNT-dependent NADPH generation is among the pathways required to maintain the various antioxidant systems in mitochondria which use NADPH, including glutathione and thioredoxin, (Chinopoulos, 2026). Thus, NNT activity dampens ROS production, as shown in brain mitochondria (Lopert and Patel, 2014; Francisco et al., 2018). The lack of MAS activity in Aralar-KO neurons limits NADH shuttling from cytosol to mitochondria reducing its capacity to regenerate NADPH. Most importantly, the lack of Aralar causes pyruvate to shift away from mitochondria toward lactate formation, thereby decreasing pyruvate-dependent mitochondrial NADH production (Pérez-Liébana et al., 2022). Taken together, these findings clearly indicate that production of mitochondrial NADH is limited in Aralar-KO neurons and this most likely jeopardises the regeneration of mitochondrial NADPH, allowing for increased ROS levels.

In agreement with these results, we now find evidence of oxidative stress in Neu Aralar-KO striatum (increased malonaldehyde reactivity). While ROS arise from the Aralar deficient neurons, our results indicate that they diffuse away into the extracellular medium and neighbouring cells, as shown by the appearance of astrogliosis, a known glial response to oxidative stress (Fulton et al., 2021; Lee et al., 2026), the upregulation of Me1 (Padj: 0.046 ; Log_2_foldchange: 0.56), a cytosolic NADPH producing enzyme enriched in glial cells (Kurz et al., 1993; McKenna et al, 1995) and induced by oxidative stress (Yang et al., 2026), and the upregulation of ALDH3B2, also enriched in astrocytes (Han et al., 2022). Remarkably, a glutathione peroxidase present in the extracellular medium, Gpx3 (Zhang et al., 2024), was the top DEG within the glutathione pathway. These findings clearly show that the intercellular milieu in the Neu Aralar-KO striatum is that of oxidative stress, to which mesencephalic DAergic terminals are particularly vulnerable (Burbulla et al., 2017; Surmeier et al., 2017). Interestingly, non-cell-autonomous mechanisms are increasingly considered to play a role in the pathogenesis of DAergic neurons (Tchung et al., 2026).

In mesencephalic presynaptic terminals from VTA, the DA transporter (DAT; SLC6A3) and vesicular monoamine transporter 2 (VMAT2; SLC18A2) coordinate DA handling by mediating its reuptake and vesicular storage, respectively (Lohr et al., 2017). Under physiological conditions, VMAT2 efficiently packages cytosolic DA into synaptic vesicles, thereby limiting its oxidation and the generation of ROS. Conversely, expansion of the cytosolic DA pool promotes DA oxidation and oxidative stress. In Neu Aralar-KO striatal terminals, the increased DOPAC/DA ratio suggests enhanced cytosolic DA turnover and a relative shift away from vesicular storage. Although VMAT2 function was not directly assessed, this phenotype may reflect an altered balance between DAT-mediated uptake and VMAT2-dependent vesicular sequestration. Consistent with the activation of compensatory mechanisms aimed at preserving DA homeostasis, Slc18a2/VMAT2 transcript levels were significantly increased in the Neu Aralar-KO striatum (Figure 3G, J). Importantly, oxidative stress has been shown to impair VMAT2-dependent vesicular DA sequestration (Fountaine et al., 2008; Cheng et al., 2008; Guo et al., 2008), whereas reduced VMAT2 activity results in decreased striatal DA content and enhanced DA turnover (Caudle et al., 2007). Therefore, increased cytosolic DA oxidation may further exacerbate the oxidative burden associated with Aralar deficiency, establishing a self-reinforcing cycle of DAergic dysfunction and ROS production within the mesolimbic pathway. Given the critical role of the VTA-to-NAc DAergic signaling in motivated social behavior (Grace et al., 2007; Gunaydin et al., 2014), these alterations may underlie the sociability deficits observed in Neu Aralar-KO mice.

### 3. β-Hydroxybutyrate restores social behavior, reduces oxidative stress, and normalizes DA metabolism in Neu Aralar-KO mice

As a major finding of this study, chronic BHB treatment restored sociability, normalized metabolic profiles, and prevented oxidative stress in the striata of Neu Aralar-KO mice. The neuroprotective actions of BHB are likely mediated by several complementary mechanisms, including the restoration of mitochondrial energy metabolism and NADH production, direct antioxidant effects, epigenetic regulation through histone deacetylase inhibition and β-hydroxybutyrylation, and possibly GPCR-mediated signaling, although the latter is thought to be primarily glial and immunomodulatory (Rahman et al., 2014).

Notably, the transcriptional profile of BHB-treated control striata closely resembled that observed in untreated Neu Aralar-KO striata, suggesting that a substantial fraction of the transcriptional response elicited by BHB in control neurons appears to be constitutively present in untreated Neu Aralar-KO neurons. KD has been reported to promote mitochondrial biogenesis and bioenergetics in neurons, through the master regulator PGC-1α (Lauritzen et al. 2016; Hasan-Olive et al., 2019). Accordingly, we previously demonstrated that BHB, the main KB produced in KD, increases PGC-1α mRNA levels in control neurons, paralleling the upregulation observed in untreated Aralar-KO neurons (Perez-Liébana et al., 2020); but failed to induce PGC1α expression in Aralar-KO ones. Moreover, whereas BHB induces antioxidant genes in control cells (Shimazu et al., 2013), untreated Neu Aralar-KO striata showed upregulation of a different set of genes associated with oxidative stress defense (Figure 4K). Therefore, Aralar-deficiency per se induces a metabolic rewiring that closely recapitulates the molecular changes typically elicited by BHB. Consistent with this notion, BHB treatment altered the expression of only 13 genes (11 upregulated) in Neu Aralar-KO mice. These findings suggest that the beneficial effects of BHB in Neu Aralar-KO mice are unlikely to arise primarily from its transcriptional actions. Rather, they are more likely to reflect the correction of key metabolic alterations associated with Aralar deficiency.

Importantly, BHB constitutes an effective substrate enabling neurons to bypass the energetic constraints imposed by Aralar-MAS deficiency. The lack of Aralar prevents adequate glucose-derived pyruvate supply to the mitochondria, inducing a metabolic limitation with deficits in mitochondrial respiration (Llorente-Folch et al., 2013a). Consistent with this, BHB effectively rescues both basal and glutamate-stimulated respiration in Aralar-deficient neurons, particularly under conditions of increased energetic demand such as neuronal hyperexcitation, a likely feature of Aralar-deficiency (Pérez-Liébana et al., 2020; Llorente-Folch et al., 2023). Once imported into mitochondria, BHB contributes to acetyl-CoA and citrate synthesis while increasing mitochondrial NADH production (Pérez-Liébana et al., 2020). In Aralar-deficient neurons, the elevated cytosolic NADH/NAD⁺ ratio (Pardo et al., 2006) drives the reduction of oxaloacetate to malate via cytosolic malate dehydrogenase to regenerate NAD⁺. BHB metabolism is likely to enhance malate-to-citrate flux by supplying substrates for the citrate/isocitrate carrier, SLC25A1, thereby supporting the operation of a citrate-malate redox shuttle. This metabolic rewiring improves mitochondrial fitness by increasing NADH availability, replenishing TCA cycle intermediates, restoring redox homeostasis, and ultimately promoting glutathione regeneration (Cavaleri and Bashar, 2018).

In conclusion, our findings reveal that neuronal Aralar deficiency disrupts social behavior through mechanisms that extend beyond the affected neurons themselves. Despite preserved Aralar expression in mesostriatal DAergic neurons, metabolic dysfunction within striatal neurons is sufficient to alter the local redox milieu and compromise DA homeostasis in the VTA terminals, identifying a novel non-cell-autonomous pathway linking neuronal bioenergetics to social-reward circuitry. Importantly, BHB effectively rescued both molecular and behavioral abnormalities, demonstrating that the consequences of Aralar deficiency remain amenable to metabolic intervention. These results establish mitochondrial metabolic dysfunction as a driver of ASD-related behavioral phenotypes and support metabolism-based therapeutic strategies for neurodevelopmental disorders associated with impaired neuronal energy homeostasis.

## MATERIALS AND METHODS

### Animals and housing

A conditional Aralar*^ff/ff^* mutant mice were generated from ES cell clones derived from C57BL/6N-A/a mice (EuMMCR; HEPD1054_A02) with loxP recombination sequences flanking exon 3 of *Aralar* gene. Targeted ES cell clones were injected into blastocysts in the transgenesis facility of CNB-CBMSO to obtain chimeric mice. These mice were crossed with a transgenic mouse expressing embryonic Flipase to eliminate the neomycin resistance selection cassette and backcrossed into C57BL/6J for eight generations (Garcia-Lobo et al., 2026). The mice obtained carry the Tm1c allele of Aralar (*Aralar^ff/ff^*). The resulting Aralar^fl/fl^ mice were crossed with CaMKIIα-Cre B6.Cg-Tg(Camk2a-cre)T29-1Stl/J; The Jackson Laboratory, stock no. 005359, RRID:IMSR_JAX:005359) and Rosa-Ai6 (B6.Cg-Gt(ROSA)26Sor^tm6(CAG-ZsGreen1)Hze/J; stock no. 007906, RRID:IMSR_JAX:007906) mouse lines to generate neuronal-specific Aralar knockout mice (Neu Aralar-KO, Aralar^fl/fl^; *CaMKIIαCre^+/-^*; *Rosa-AiC^+/-^*) and control littermates. Adult male and female mice of C57BL/6J background were used, maintained on a 12-hour light/dark cycle with food and water provided ad libitum. Unless otherwise specified, both sexes were included in all tests.

All animal experimentation procedures conformed to the European Guidelines for the Care and Use of Laboratory Animals (Directive 86/609) and were approved by the Ethical Committees for Animal Experimentation of the Centro de Biología Molecular Severo Ochoa, Universidad Autónoma de Madrid and the Comunidad Autónoma de Madrid (PROEX 305.0/21), Spain. Reporting followed the ARRIVE Guidelines. All efforts were made to minimize the number of animals used and their suffering.

### Drug administration

In vivo β-hydroxybutyrate (BHB) administration was performed by intraperitoneal injection of 520 mg/kg/day dissolved in 0.9% NaCl for 15 consecutive days in 7–9-month-old female Neu Aralar-KO mice and controls. For evaluation of tyrosine hydroxylase activity, the aromatic L-amino acid decarboxylase (AADC) inhibitor NSD1015 (Sigma-Aldrich, #54880) was injected intraperitoneally at 100 mg/kg in 0.9% NaCl 1 hour before sacrifice in 7–9-month-old Neu Aralar-KO and control mice.

### Behavioral analysis

Adult mice (7–9 months old) were acclimated for 7 days in the behavioral facility and handled daily (5 min/day) for 3 days before testing. All behavioral experiments were conducted between 10:00 and 18:00 during the light phase, with animals habituated to the testing room for 30 minutes prior to trials. Experiments proceeded from the least to most stressful tests. Animals were tested individually, and apparatuses were cleaned with 70% ethanol between trials to eliminate odor cues. Experimenters were blinded to genotype.

### Open Field Test

Mice were placed in a 40 × 40 × 35 cm open field arena with transparent walls and a black plastic bottom lined with fresh wood shavings for each mouse. Illumination was provided by a 25 W halogen bulb positioned 180 cm above the arena. Exploration behavior was recorded for 10 minutes. The ANY-maze software (Stoelting®) was used to analyze movements via a virtual grid of 10 × 10 cm squares (Krauter et al., 2019a).

### Y-maze Test

Spatial memory was assessed using a Y-maze with three 50 cm arms arranged at 120° angles and 20 cm high black walls (Kraeuter et al., 2019b). Arms were randomly assigned as start (open), novel (closed during training, open during testing), and other (open). The test consisted of two trials 24 hours apart. During the 10-minute training trial, mice explored two arms (start and other) while the novel arm was blocked. In the 5-minute test trial, all arms were accessible. Entries and time spent in each arm were recorded.

### Novel Object Recognition (NOR) test

Episodic-like memory was evaluated in the open field arena by exploiting the rodents’ preference for novelty (Leger et al., 2013). During training, mice explored two identical objects (chess pieces) for 10 minutes. After 24 hours, mice were reintroduced to the arena containing one familiar and one novel object placed in the same location. Exploration was defined as sniffing or touching with the nose/forepaws. ANY-maze software quantified exploration time. Recognition performance was expressed as a relative discrimination index (rDI) calculated as (T_N_ – T_F_)/(T_N_ + T_F_), where T_N_ and T_F_ represent time exploring the novel and familiar objects, respectively.

### Three-chamber social behavior test

A transparent acrylic box divided into three interconnected chambers was used to assess social interaction and preference (Yang et al., 2011). Mice (7–9 months, females) were habituated for 5 minutes with two empty wire cages in the outer chambers. Twenty-four hours later, an unfamiliar age-and sex-matched conspecific was placed under one cage and an inanimate object under the other. The subject mouse explored freely for 7 minutes. Time spent sniffing within proximity to the cages was recorded. Social preference and memory were quantified using Social preference index calculated as T_S_/(T_S_+T_O_), where T_S_ and T_O_ represent the time exploring the conspecific and inanimate objects respectively. Stimulus mice were habituated to the wire cages for 15 min before testing.

### Morris Water Maze (MWM)

Spatial learning and memory were evaluated in a circular pool (120 cm diameter) filled with water at 22–24°C and a submerged escape platform (10 cm diameter) fixed during acquisition (Vorhees et al., 2006). Visual cues were placed around the room. On day 0, mice were habituated with a 60-second swim without the platform. Acquisition was conducted over three days (days 1–3) with three trials/day from pseudo-random start positions (up to 60 seconds each). Failure to locate the platform resulted in guidance to it for 10 seconds. Escape latency, measured as time spent finding the platform since release was recorded. On day 4, a 60-second probe trial tested memory with the platform removed; time spent in the target quadrant and platform crossings were recorded. For reversal learning (days 5–7) platform was relocated to the opposite quadrant and escape latency was recorded again as mice tried to find the novel position of the platform.

### Dendritic spine labeling

Coronal brain slices (300 μm) were prepared in oxygenated artificial cerebrospinal fluid (aCSF) (119mM NaCl, 2.5mM KCl, 1 NaH₂PO₄, 11mM glucose, 26mM NaHCO₃, 1.2mM MgCl₂ and 2.5mM CaCl₂) and allowed to recover at 32°C for 1 hour. Slices were labeled with 10 μM DiI-coated glass micropipettes, incubated for an additional hour, fixed in 4% paraformaldehyde for 10 minutes, washed, and mounted. Confocal images were acquired using a Zeiss LSM800 microscope (100×/1.4 NA oil immersion objective) (Foster et al., 2017). Spines were manually counted and classified using FIJI software from three fields per slice across three brain sections per animal.

### HPLC analysis of amino acids and monoamines

Brains were rapidly removed and cut into 1 mm coronal slices on ice using a coronal rodent brain matrix (Electron Microscopy Sciences, Hatfield, PA, USA). The ventral striatum was dissected and stored at −80 °C. Samples were sonicated in 6 volumes (w/v) of 0.4 N perchloric acid for deproteinization and centrifuged at 10,000 × g for 20 min at 4 °C (Pérez-Liébana et al., 2020). Amino acids in the supernatant were analyzed by HPLC with fluorescence detection (Jasco FP-2020; λex = 240 nm, λem = 450 nm) as previously described (Perucho et al., 2015). Monoamines and their metabolites were analyzed in the same supernatants by HPLC with an ESA Coulochem electrochemical detector as described (Mena et al. 1984) with minor modifications. Separation was performed on an ACE 5 C18 column (150 × 4.6 mm; Advanced Chromatography Technologies, Aberdeen, UK). The mobile phase was 0.1 M citrate/acetate buffer, pH 3.9, containing 10% methanol, 1 mM EDTA and 1.2 mM heptane sulfonic acid, at 1 mL/min. Detector potentials were +0.40 V (guard cell), +0.05 V (D1) and −0.39 V (D2). Compounds were identified by retention time and quantified against external standards (0.6 µM). Results are expressed as nanogram of metabolite/gram of tissue.

### RNA sequencing and transcriptomic analysis

Total RNA was extracted from frozen ventral striatal tissues dissected using a coronal rodent brain matrix (Electron Microscopy Sciences, Hatfield, PA, USA) (slice interval1 mm) using the Maxwell® RSC SimplyRNA Tissue Kit on a Maxwell® RSC 48 Instrument (Promega), following the manufacturer’s instructions. RNA integrity was assessed (RIN > 7, Agilent 2100). Sequencing was performed by Novogene (Cambridge, UK). Strand-specific libraries were prepared after ribosomal RNA depletion. Libraries were then sequenced on an Illumina NovaSeq X Plus Series (paired-end, 2 x 150bp), yielding 60.7-83.1 million read pairs per sample.

Adapter-containing and low-quality reads were removed and read quality was assessed with FastǪC (v0.12.1) and MultiǪC (v1.18). Clean reads were aligned to the mouse reference genome GRCm39 (Ensembl release 113) using HISAT2 (v2.1.0), with 88.8–94.3% of reads aligned per sample. Reads were assigned to genes with htseq-count (v0.11.2), using reverse-stranded, intersection-strict mode. Because the experimental groups were not balanced for sex, Y-chromosome genes were removed from the annotation before counting.

Differential expression analysis was performed in R (v4.2.2) with DESeq2 (v1.38.2). The design included four groups: Control (n = 3), Neu-Aralar-KO (n = 3), Control + BHB (n = 4) and Neu-Aralar-KO + BHB (n = 6), n being the number of individuals. Pairwise comparisons were made. P values from Wald tests were adjusted using the Benjamini–2Hochberg procedure. Genes with an adjusted p < 0.05 and |log2 fold change| > 1 were considered differentially expressed. Genes were annotated using Ensembl BioMart via biomaRt. Gene Set Enrichment Analysis (GSEA) was performed on genes ranked by the DESeq2 test statistic against the Molecular Signatures Database (MSigDB). Gene Ontology over-representation analysis of DEGs was performed with topGO.

### Metaplot analysis of H3KGbhb enrichment

Publicly available ChIP-seq data for histone H3 lysine 9 β-hydroxybutyrylation (H3K9bhb) in the cerebral cortex of C57BL/6J mice subjected to 48 h of fasting, relative to ad libitum-fed controls, were obtained from GEO (accession GSE168723; Cornuti et al., 2023). Fasting-enriched ChIP-seq peaks, originally mapped to the mm10 genome assembly, were converted to mm39 coordinates using the UCSC LiftOver tool. The resulting bedGraph file was converted to bigWig format using bedGraphToBigWig. deepTools plotProfile was used to generate metaplots of H3K9bhb signal across the gene bodies (TSS to TTS) of upregulated differentially expressed genes (DEGs) identified by RNA-seq in this study, using a bin size of 50 bp. As a control, the corresponding gene-body regions were randomly shuffled across the genome using bedtools shuffle, and H3K9bhb signal was profiled across these shuffled regions using the same parameters.

## Acknowledgements

We thank Bárbara Sesé for excellent technical assistance; Belén Pintado and Verónica Dominguez from CNB Transgenesis unit; Rosa Sedano from SIdI-UAM facility; the Advanced light microscopy (SMOA) and Animal facilities of the CBMSO for their support. The *Rosa-AiC* (B6.Cg-Gt(ROSA)26Sortm6(CAG-ZsGreen1)Hze/J) mouse line was kindly provided by Dr. Javier Palazuelos (UCM, Madrid, Spain). The *CaMKIIα-Cre* (B6.Cg-Tg(Camk2a-cre)T29-1Stl/J) mouse line was kindly provided by Dr. JM Cuezva (CBMSO, UAM-CSIC, Madrid, Spain) and Dr. Teresa Iglesias (IIB, UAM-CSIC, Madrid, Spain).

## Supplementary Material

### Supplementary Materials and Methods

#### Genotyping

Mouse genotypes were determined by PCR from tail or ear DNA extracted by incubation in 50 mM NaOH at 95°C for 30 minutes, neutralized with 1 M Tris-HCl pH 7.5, and centrifuged (14,000 rpm, 5 minutes). PCR amplification followed standard protocols. For Cre Recombinase genotyping we used the following oligonucleotides: Cre_Fw 5’-CATTTGGGCCAGCTAAACAT-3’ and Cre_Rv 5’-TAAGCAATCCCCAGAAATGC-3’. For Aralar^fl/fl^ we used the following oligonucleotides: Aralar^fl/fl^_Fw 5’-TGGATATTTTGCCTGCACTTGTCTC-3’ and aralar^fl/fl^_Rv 5’-CCTGAAAATCTGCTGGGTGAAATG-3’. For ROSA:Ai6 we used the following oligonucleotides: Rosa Mut_Fw 5’-GGCATTAAAGCAGCGTATCC-3’ and Rosa Mut_Rv 5’-AACCAGAAGTGGCACCTGAC-3’.

#### In vivo MRI and ¹H MRS

Mice were anesthetized with isoflurane (2% in 1 L/min O₂) and maintained at 70 ± 20 breaths/min and ∼37 °C (SA Instruments monitoring system). Imaging was performed on a 7.0 T Bruker BioSpec with a 23 mm ¹H volume coil and ParaVision 6.1. Axial T₂-weighted RARE images were acquired with TR/TE = 2500/26 ms, RARE factor 8, 6 averages, FOV 23 × 23 mm², matrix 256 × 256, and 14 contiguous 1 mm slices (Hennig et al., 1986). Ventricular volume was calculated as the sum of lateral ventricle areas across slices multiplied by slice thickness, using the Paxinos and Franklin atlas as reference. ¹H MRS was acquired from a [3 × 3 × 3 mm³] voxel in the right cortex, hippocampus and thalamus. It used PRESS (Bottomley, 1987) with VAPOR water suppression (Tkáč et al., 1999), TR/TE = 2500/35 ms and 128 averages. Spectra were quantified with LCModel [v6.2-0R] (Provencher, 1993).

#### Western blot

Cortex, hippocampus and striatum were dissected using a coronal rodent brain matrix (Electron Microscopy Sciences, Hatfield, PA, USA) (slice interval1 mm). Tissue was homogenized in RIPA buffer containing the protease inhibitors phenylmethylsulfonyl fluoride (PMSF, 1 mM; Fluka) and iodoacetamide (1 mM; Merck). Lysates were sonicated and centrifuged [14,000 × g, 15 min, 4 °C], and protein concentration in the supernatant was determined by BCA assay. Proteins (20 µg) were separated on 8% SDS-PAGE gels and transferred to nitrocellulose membranes. Membranes were blocked in 5% non-fat milk or BSA and incubated overnight at 4 °C with primary antibodies (anti-Aralar (del Arco and Satrústegui, 1998),1:4000; anti-MAG, Santa Cruz, Sc166849,1:1000; anti-Tubulina, Sigma-Aldrich, T5168, 1:5000). They were then incubated with HRP-or Alexa Fluor-conjugated secondary antibodies. Signals were detected by chemiluminescence or fluorescence and quantified by densitometry in ImageJ. Malondialdehyde (MDA)–protein adducts were detected with an anti-MDA antibody (1:1000; Abcam, ab27642). The MDA signal was quantified over the entire lane and normalized to total protein stained with Ponceau S (Giera et al., 2012; Dharmajaya et al., 2022).

#### Immunohistochemistry

For immunohistochemistry, sections were abundantly washed (1 X PBS) and treated with NaBH4 (1 mg/ml in PBS pH 8.0) to quench endogenous autofluorescence. Free-floating sections were blocked for 1 h in PBS containing 10% normal horse serum, 0.25 % Triton X-100 and incubated O/N at 4°C with primary antibodies (anti-MBP, Biorad, MCA685S, 1:300; anti-TH, Pel-Freeze,1:300; anti-GFAP, Dako, Z0334, 1:300). Afterwards, sections were rinsed three times in PBS containing 0.25 % Triton X-100 and then incubated for 2 h with the secondary antibodies conjugated with different alexa fluorophores. After three washes with PBS containing 0.25 % Triton X-100, sections were stained for nuclei using either DAPI (Merck ® #268298) for 10 minutes at 1:10000 dilution. Sections were mounted onto poli-Lysine-coated slides, dehydrated, and coverslipped with Mowiol (Dabco®).

Images were acquired using a Nikon A1R+ confocal inverted microscope. Nikon 20X/0.75 Plan-Apo air objective (MRD00205) was employed. Image acquisition was carried out with NIS-Elements software (v. 5.30.02). DAPI was excited using 405 diode laser, Alexa 550 was excited using 561 diode laser and Alexa 647 was excited using a 640-diode laser. Signals were sequentially scanned using 1024x1024 scanning area producing a 0.15 um pixel size. Tile scans shown generated by stitching 6 × 6 fields in NIS-Elements with 15% overlap, using blending mode and a fully motorized stage. All imaging was performed at the Advanced Light Microscopy Facility, Centro de Biología Molecular Severo Ochoa (Madrid, Spain).

## Supplementary figures

**Supplementary Figure S1.**
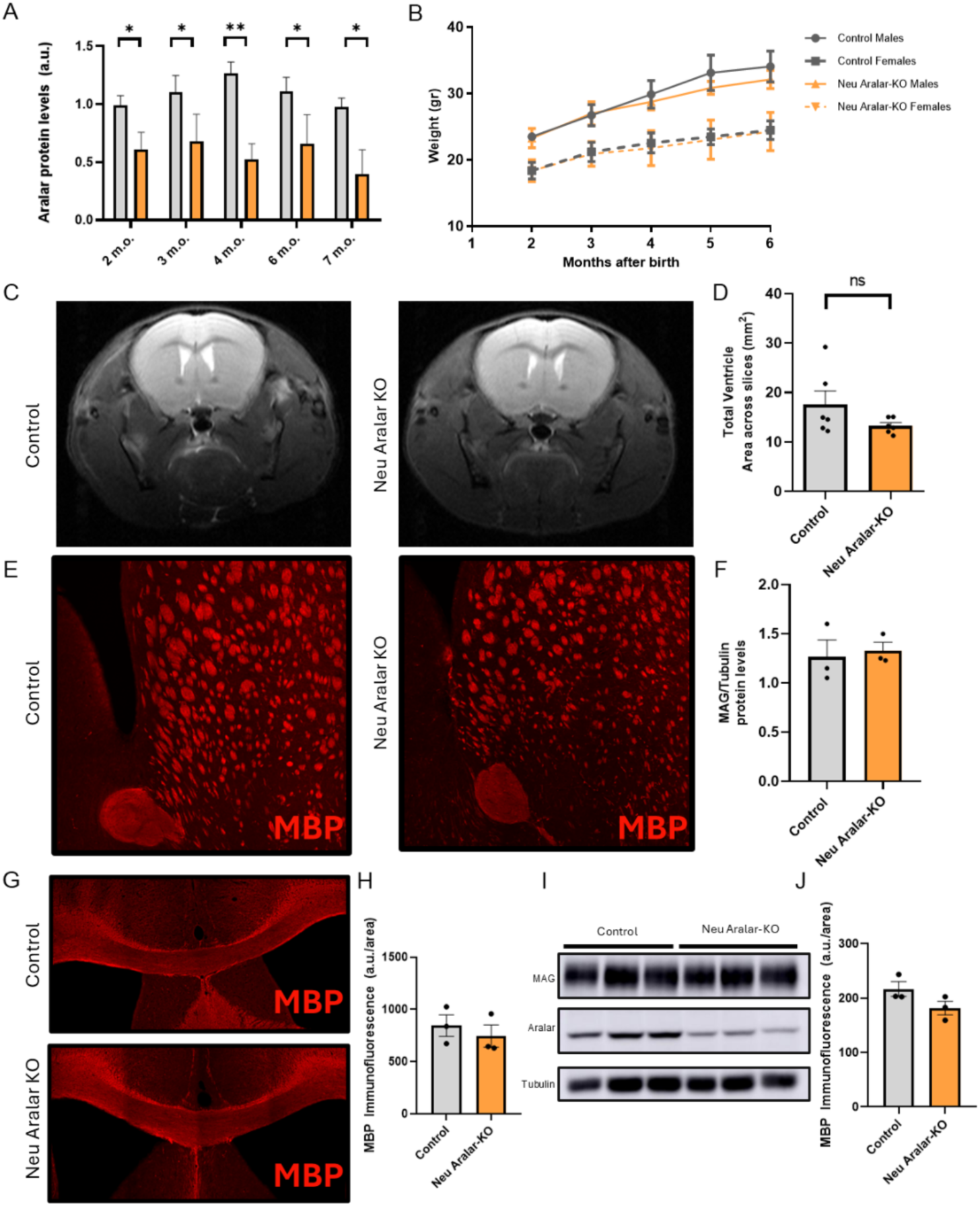
Characterisation of the Neu Aralar-KO model: Aralar loss over time, growth, brain morphology and myelination. (A) Ǫuantification of Aralar protein levels in Control and Neu Aralar-KO mice at 2, 3, 4, 6 and 7 months of age (m.o.). (B) Body weight of Control and Neu Aralar-KO males and females from 2 to 6 months after birth. (C) Representative T2-weighted magnetic resonance images of Control (left) and Neu Aralar-KO (right) brains. (D) Ǫuantification of total ventricular area across slices. (E) Representative images of myelin basic protein (MBP) immunostaining (red) in the striatum of Control (left) and Neu Aralar-KO (right) mice. (F) Ǫuantification of MAG protein levels normalised to tubulin. (G) Representative images of MBP immunostaining (red) in the corpus callosum of Control (top) and Neu Aralar-KO (bottom) mice. (H) Ǫuantification of MBP immunofluorescence in the corpus callosum (arbitrary units per area). (I) Representative immunoblot of MAG, Aralar and tubulin in Control and Neu Aralar-KO samples. (J) Ǫuantification of MBP immunofluorescence in the striatum (arbitrary units per area). Data are presented as mean ± SEM (n = 3 mice per group except for RMN where n= 6 mice per group). Statistical analysis was performed using unpaired Student’s *t*-test; n.s., not significant; *p ≤ 0.05, **p ≤ 0.01. Scale bars, 500 µm (E, G).

**Supplementary Figure S2.**
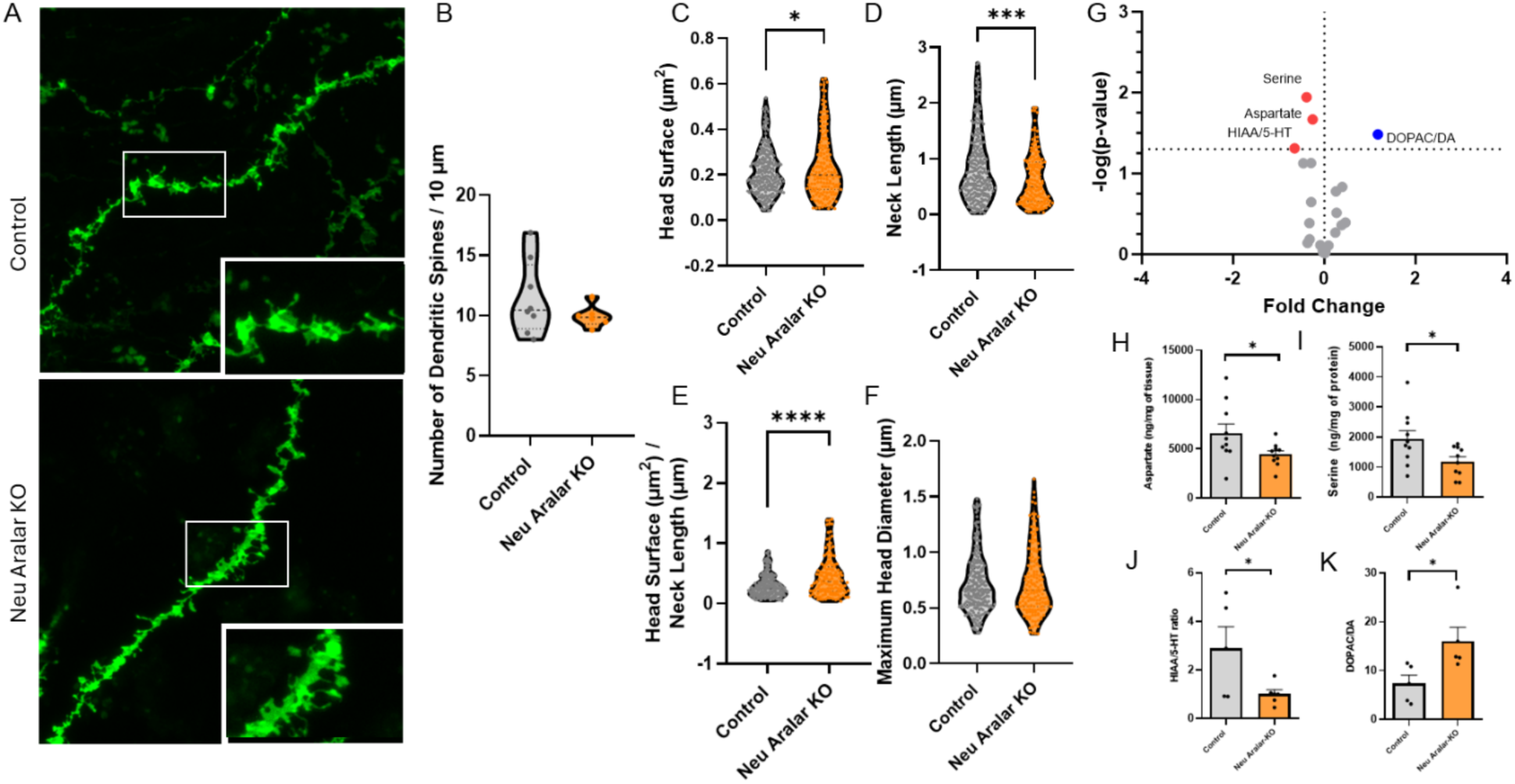
Dendritic spine morphology and metabolite levels in the cortex of Neu Aralar-KO mice. (**A**) Representative images of dendritic spines in cortical neurons from Control (top) and Neu Aralar-KO (bottom) mice following DiI microcrystal labelling; insets show the boxed regions at higher magnification. Ǫuantification of (**B**) dendritic spine density, (**C**) head surface area, (**D**) neck length, (**E**) head surface area-to-neck length ratio and (**F**) maximum spine head diameter. (**G**) Volcano plot of amino acid and monoamine alterations in the cortex of Neu Aralar-KO relative to Control mice; significantly changed metabolites are coloured and labelled, and the dotted horizontal line indicates the significance threshold. Ǫuantification of (**H**) aspartate, (**I**) serine, (**J**) the 5-HIAA/5-HT ratio and (**K**) the DOPAC/dopamine ratio in cortex. Data are presented as mean ± SEM (n = 5 mice per group). Statistical analysis was performed using unpaired Student’s *t*-test; n.s., not significant; # denotes a non-significant trend; *p ≤ 0.05, ***p ≤ 0.001, ****p≤ 0.0001.

**Supplementary Figure S3.**
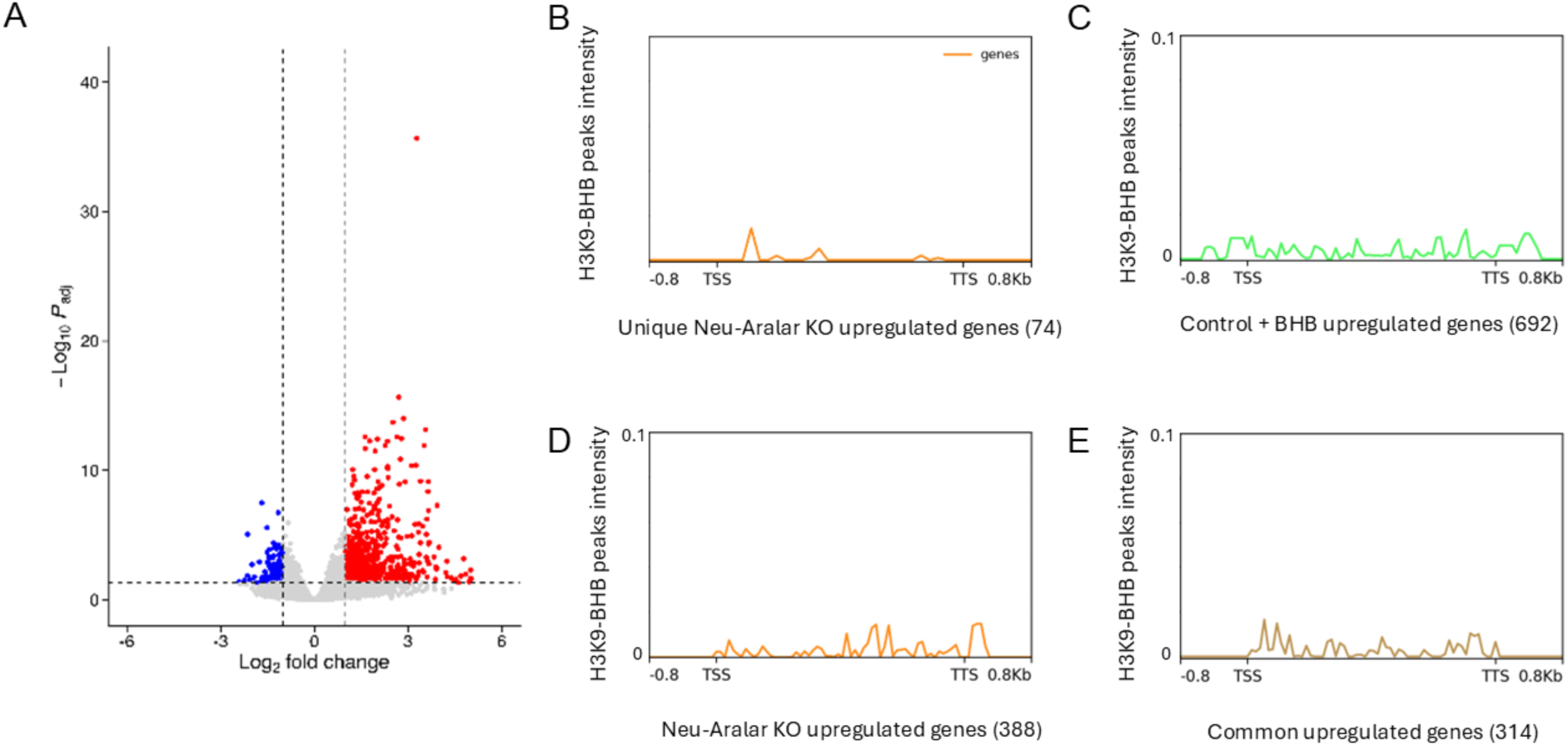
Transcriptional response to β-hydroxybutyrate and its relationship with H3KG-β-hydroxybutyrylation. (**A**) Volcano plot showing differentially expressed genes in BHB-treated Control mice relative to untreated Control mice. Blue dots indicate significantly downregulated genes, red dots significantly upregulated genes and grey dots non-significant genes (adjusted p < 0.05, |log_2_FC| > 1). (**B–E**) Metaplots of H3K9-bhb ChIP-seq signal intensity (Cornuti et al., 2023) over (**B**) the 74 genes upregulated uniquely in the Neu Aralar-KO *versus* Control comparison, (**C**) the 388 genes upregulated in Neu Aralar-KO *versus* Control, (**D**) the 692 genes upregulated in Control + BHB *versus* Control and (**E**) the 314 genes common to both comparisons. Signal is displayed from 0.8 kb upstream of the transcription start site (TSS) to 0.8 kb downstream of the transcription termination site (TTS).

